# Carotenoid-based ornaments predict survival in male but not female birds: a meta-analysis

**DOI:** 10.64898/2026.01.21.700769

**Authors:** C Alonso-Alvarez, M Briga, J Morales, AA Romero-Haro

## Abstract

Many vertebrates exhibit colourful ornaments generated by carotenoid pigments that presumably evolve as signals under sexual selection. To function as sexual signals, carotenoid-based coloured ornaments should accurately reveal an individual’s quality, commonly defined as fitness potential, which includes the ability to survive. We performed a meta-analysis for testing the association between the expression level of carotenoid-based ornaments and longevity estimates (annual survival or lifespan) in birds, the taxon with the largest body of evidence. We detected a significant positive correlation in males but not in females. This result contrasts with recent meta-analytic work showing a positive association between carotenoid-based coloured traits and avian fecundity in both sexes. The link to survival was consistent among colourations produced by enzymatically transformed pigments (mostly red ketocarotenoids) and those based on dietary yellow carotenoids. The latter suggests that the association with survival holds across the physiological mechanisms involved in the production of these ornaments. In summary, our study reveals sex-specific selection in the evolution of carotenoid-based ornaments.

## 1. INTRODUCTION

Among animals, informative signals are traits that have evolved to communicate information that benefits both the emitter and receiver (e.g., Maynard Smith and Harper 2004; Penn and Szamado 2020). Potential benefits include the emitter gaining access to resources or mating opportunities, while the receiver may avoid social conflicts or attain high-quality mates (Maynard Smith and Harper 2004). In a sexual selection context, high-quality mates can provide either more direct benefits, such as resources or care for reproduction (the “good parents” hypothesis; Hoelzer 1989), or indirect ones, such as the transmission of genes to descendants that make offspring more sexually attractive (“sexy-son” hypothesis; Weatherhead and Robertson 1979) or more viable (“good genes” hypothesis; Hamilton and Zuk 1982).

The acquisition of these benefits depends on how reliably the signal indicates the bearer’s quality (Maynard Smith and Harper 2004; Wilgers and Hebets 2015). Traditionally, researchers have attempted to demonstrate signal reliability by correlating signal intensity or size with the signaller’s potential for survival and fecundity (i.e., potential fitness). Thus, a vast body of research has investigated the association between signal expression and measures of individual condition, such as size-corrected body mass, physiological and oxidative stress levels, immunocompetence or parasite resistance (e.g., see meta-analyses in Guindre-Parker and Love 2014; Moore et al. 2016; Weaver et al. 2018; White et al. 2020; Dougherty 2021). However, individual condition is an elusive concept often used as a fitness proxy (Clancey and Byers 2014; Ronget et al. 2017). Instead, other meta-analyses have detected associations between the expression levels of sexual ornaments and fecundity-related parameters (Weaver et al. 2018; Hernández et al. 2021; Nolazco et al. 2022). However, establishing a direct link with survival is more challenging because this fitness component must be estimated from longitudinal monitoring (e.g., survival rates or longevity records), which often entails substantial logistical constraints. Consequently, studies adopting this approach are scarce compared to those focusing on condition or fecundity (see Jennions et al. 2001; Nolazco et al. 2022; Pollo et al. 2025).

Among ornaments involved in sexual signalling, the conspicuous colourations generated by carotenoid pigments (yellow to red traits) have attracted the attention of evolutionary biologists since Darwin and Wallace (e.g., Caro 2017). Exploring the proximate mechanisms underlying the production of carotenoid-based colourations has enhanced our understanding of how ornaments become reliable signals (e.g., Olsson and Owens 1998; Hartley and Kennery 2004; Hill et al. 2023). Carotenoids must be acquired along with food, meaning they may constitute a limited resource under natural conditions (Endler 1980; Kodric-Brown 1985; Hill 1990; but see also Hadfield and Owens 2006 or Simons et al. 2014). They are also considered antioxidants (von Schantz et al. 1999; Pérez-Rodríguez 2009; Svensson and Wong 2011; Nabi et al. 2020; but see also Koch et al. 2018). It has therefore been hypothesized that a resource allocation trade-off between investing in these pigments for colouration and homeostasis mediates signal reliability, with lower-quality individuals incurring a higher fitness cost when expressing such signals (e.g., Møller et al. 2000; Alonso-Alvarez et al. 2008). Indeed, a meta-analysis showed a positive correlation between carotenoid-based colourations in birds and T-cell-mediated immune response and between carotenoid levels and antioxidant status (Simons et al. 2012a), at least partially supporting the role of carotenoids in homeostasis and, therefore, potentially in survival or longevity (e.g., Pike et al. 2007; Freeman-Gallant et al. 2011).

A second mechanism may also sustain the reliability of some carotenoid-based ornaments. Many red and some yellow ornaments are produced from carotenoids that are synthesized endogenously from yellow dietary carotenoids (McGraw 2006). It has been suggested that enzymes closely related to cellular respiration perform this carotenoid transformation. Hence, the signal expression level may directly reflect individual quality without involving expression costs or resource-allocation trade-offs (i.e., the "shared-pathway hypothesis"; Hill 2011; Hill et al. 2023). Indeed, a meta-analysis on passerines showed a positive correlation between plumage colour intensity and parasite resistance, as well as specific reproductive parameters, but only in species that produce colourations using transformed carotenoids (Weaver et al. 2018). These authors suggested that transformed carotenoids better reflect individual quality due to the physiological links between cellular function and carotenoid metabolism, making these ornaments more reliable indicators of quality (Hill et al. 2023). However, they did not test a direct link between sexual ornaments and survival.

Sexual ornaments are often more conspicuous in males. This has traditionally been interpreted as a consequence of stronger sexual selection pressure on this sex, female-biased investment in reproduction (due to anisogamy) or higher reproductive or survival costs in females, which constrain signal evolution in this sex (see seminal works by Bateman 1948; Trivers 1972; Lande 1980). Bateman (1948) proposed that females would obtain fitness gains primarily through increased investment in self-maintenance (longevity), while males would achieve fitness gains by acquiring multiple mates through investment in sexual signalling. Other authors have suggested that, at least in some species, the presence of ornaments in both sexes may merely be due to genetic correlations (i.e., Lande 1980; Dale et al. 2015; see also Doutrelant et al. 2020 and references therein). However, the last decades have witnessed an increased interest in the evolution of female sexual signalling, which might favour a paradigm shift (Amundsen 2000; Nordeide et al. 2013; Doutrelant et al. 2020). Examples and theoretical support for the idea that female ornaments can be sustained through direct sexual or social selection (i.e., only related to resource acquisition, not mates) have grown substantially (e.g., Dale et al. 2015; Hare and Simmons 2019; Doutrelant et al. 2020). Thus, it has been hypothesized that a positive correlation between sexual signal expression and fitness in females could be the result of direct selection on this sex, whereas a negative correlation could be a consequence of genetic correlation due to sexual conflict and costs associated with the production of the female ornament (Clutton-Brock 2009; Nordeide et al. 2013; Fargevieille et al. 2023). Indeed, a meta-analysis performed exclusively on female birds detected a positive link between the expression level of carotenoid-based ornaments on bare body parts (as opposed to feathers) and clutch size (Hernández et al. 2021). Furthermore, Weaver et al. (2018) on passerine carotenoid-based plumage colourations and Nolazco et al. (2022) on a broader range of avian visual ornaments found a positive relationship between trait expression level and fecundity-linked parameters in both sexes, thus supporting the idea that direct sexual selection also promotes the evolution of female ornaments.

Despite evidence that carotenoid-based sexual signals are positively associated with reproductive success in both sexes, whether similar associations exist with survival or longevity has rarely been tested using meta-analytic approaches (Nolazco et al. 2022; Pollo et al. 2025). Here, we performed a systematic literature search and a meta-analysis to test the link between carotenoid-based ornamentation and survival in any animal species. First, we tested the hypothesis that ornaments (putative sexual signals sensu Pollo et al. 2025) coloured by carotenoid pigments predict individual survival or longevity, thereby informing potential signal receivers about a key component of the signaller’s fitness potential. If direct selection favours signalling honest information, and if the production mechanism of carotenoid-based ornaments is linked to homeostasis, we predict a positive correlation with survival or longevity in both sexes. Alternatively, females could show no link or a negative correlation if selection on signal reliability is weaker in this sex. Second, we tested Hill’s (2011) hypothesis that colourations produced by transformed carotenoids are more closely linked to individual quality than those produced by dietary pigments. Hence, we accounted for differences in sexual signalling production by testing whether ornaments based on transformed carotenoids reveal a stronger link to survival or longevity than those produced by untransformed carotenoids.

## 2. Material and Methods

### 2.1 Literature selection

We reviewed the literature to identify studies in any animal species that tested for correlations between the conspicuousness (colour intensity or size of the coloured area) of carotenoid-dependent colourations and individual survival rates or longevity. The Web of Science (WOS) library was used, which avoided non-peer-reviewed literature. The following Boolean terms were used: caroten* and (colour* or color* or bright*) and ("lifespan" or "life span" or longevity or surviv* or mortalit*) and ("trade-off*" or evolutionary or "life histor*" or signal* or "sexual selection" or behav* or "mate choice" or "individual quality" or honest or handicap). The search yielded 409 papers up until May 7^th^ 2025 (see PRISMA in Figure S1). Additionally, we reviewed the Scopus, PubMed, and Google Scholar databases, which yielded no additional relevant peer-reviewed papers. Our WOS revision reported studies in 25 animal species, including two insects (i.e., *Orgyia antiqua* in Sandre et al. 2007, and *Harmonia axyridis* in Sun et al. 2018) and two fish species (i.e., *Poecilia reticulata* in Gordon et al. 2015 and Gasparini et al. 2019, and *Gasterosteus aculeatus* in Pike et al. 2007 and Simons et al. 2021). All other studies analysed birds. Accordingly, we focused our statistical analyses on avian species to avoid phylogenetic imbalance (see below). Three avian studies were excluded as they analysed nestling carotenoid-based colourations that disappear or change in adulthood, making sexual selection infeasible (i.e., Fitze and Tschirren 2006; Pirrello et al. 2017; Enbody et al. 2021). Thus, we focused on 18 species belonging to 10 avian families (Table S1): Phasianidae (2 species), Picidae (1), Maluridae (1), Turdidae (1), Paridae (2), Estrildidae (1), Fringillidae (5), Parulidae (3), Pelecanidae (1) and Passerellidae (1).

Monitoring times varied from weeks to years, which were statistically considered (below). Each colouration subject of analysis was assigned to one of two categories: (1) colourations produced by transformed carotenoids, including both red ketocarotenoids and yellow canary xanthophylls and (2) ornaments coloured only by non-transformed dietary carotenoids (e.g., lutein, zeaxanthin, β-carotene; e.g., McGraw, 2006; Weaver et al. 2018). Identifying these carotenoid types was based on species-specific literature (see Electronic Supplementary Material). Alternatively, the colourations were classified as red (i.e., assumedly more conspicuous and produced by ketocarotenoids) or yellow traits (i.e., made by canary xanthophylls or dietary carotenoids).

### 2.2 Effect size estimation

We extracted effect size estimates (Figure 1) following Nakagawa and Cuthill (2007) and Peters et al. (2019). For all studies, we extracted either the means and variance estimates (colour intensity or ornament size) of survivors and non-survivors or test statistics such as *t* or *F* values, (sign of) *β* values, and *χ*^2^ values and converted these statistics into a standardized *Zr* value following the equations in (Lipsey and Wilson 2001) using the online calculator on the Campbell Collaboration website (https://www.campbellcollaboration.org/calculator/r-r). A positive *Zr* value means that individuals with more intense colourations or bigger coloured ornaments survive better, while a negative value indicates that individuals with less intense colouration or smaller coloured traits survive better. The sign of the relationship between the survival estimates and colour metrics was carefully considered when extracting effect sizes, e.g., hue is negatively correlated with trait redness (McGraw 2006; García de Blas et al. 2016; Cantarero et al. 2019), such that positive effect sizes consistently indicate increased expression-level and survival.

**Figure 1.**
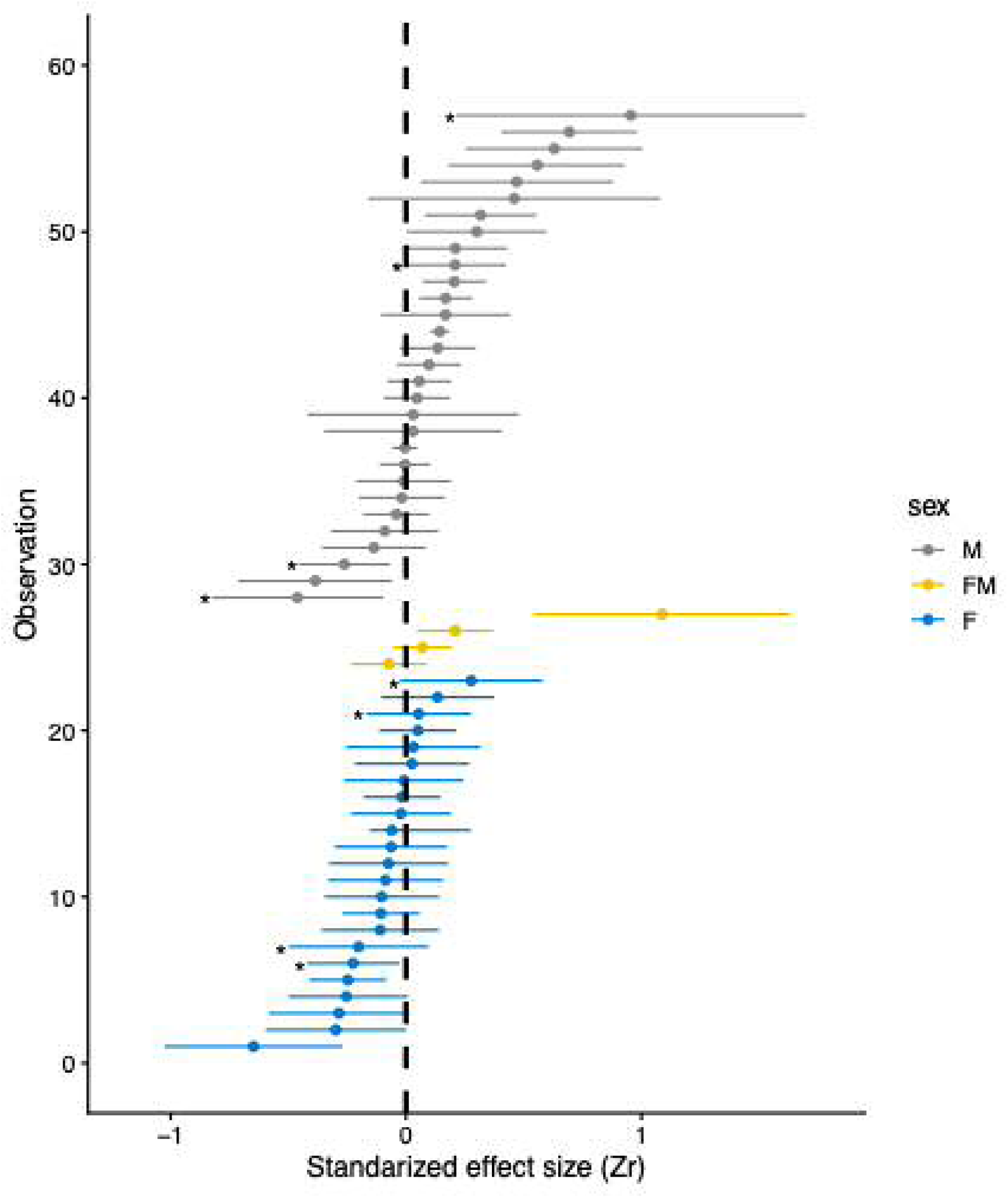
Sex-specific distribution of the 57 effect sizes and their 95% confidence intervals. Effect sizes with stars on the left represent quadratic associations and a figure without these data is shown in Electronic Supplementary Material (Figure S3). Abbreviations: M = males, F = females, FM = females and males combined.

Results reported only as delta AIC values were converted to *χ*^2^ values, adjusting the statistic for twice the difference in the number of estimated parameters. For studies that reported Cox proportional hazards, we converted the test statistic into a *χ*^2^. For studies that reported both univariate and multivariate statistics, we selected the multivariate results. When we had access to survival data, we took the survival probabilities at the longest monitoring time. For studies that reported the means and their variance in the form of graphs, we extracted values using the software WebPlotDigitizer version 5.2 (Rohatgi 2024), available at https://automeris.io/). When the results for control individuals were separated from those undergoing an experimental treatment, we selected the control group to avoid any confounds or bias arising from the experiment. We used the sample size reported with the study’s test statistic, and when no test statistic was available, we used the number of individuals. When possible, we separated the survival measurements made for first-year individuals and adults. For studies with annual monitoring, the monitoring time is the maximum number of years an individual could be recaptured.

### 2.3 Phylogenetic meta-regression

We performed a meta-regression testing for the fixed factors sex (males vs females), colour type (yellow or red) and carotenoid type (dietary untransformed vs transformed). In addition, we included several moderator variables such as the study environment (captivity vs wild), age at colour sampling (yearlings, adult or both), species maximum lifespan and study length (both in years) and the type of coloured trait (bare part vs. plumage; Table S1).

We performed the meta-regression using the techniques employed in previous studies (Briga et al. 2012; Kärkkäinen et al. 2022). In brief, we accounted for multiple measurements of the same species by including species identity as a random intercept. We accounted for the non-independence of species by including phylogeny (Figure 2) as a random term in the models’ variance-covariance matrix (Hadfield and Nakagawa 2010, de Villemereuil and Nakagawa 2014). Hence, the model contains both species and phylogeny as random intercepts. Including other random intercepts, such as study, is not possible because most species have only one study (26 studies on 18 species, Figure 2, Table S1), making both random intercepts indistinguishable. We coded residual variance explicitly as an observation-level random intercept (de Villemereuil and Nakagawa 2014; Bürkner 2017).

**Figure 2.**
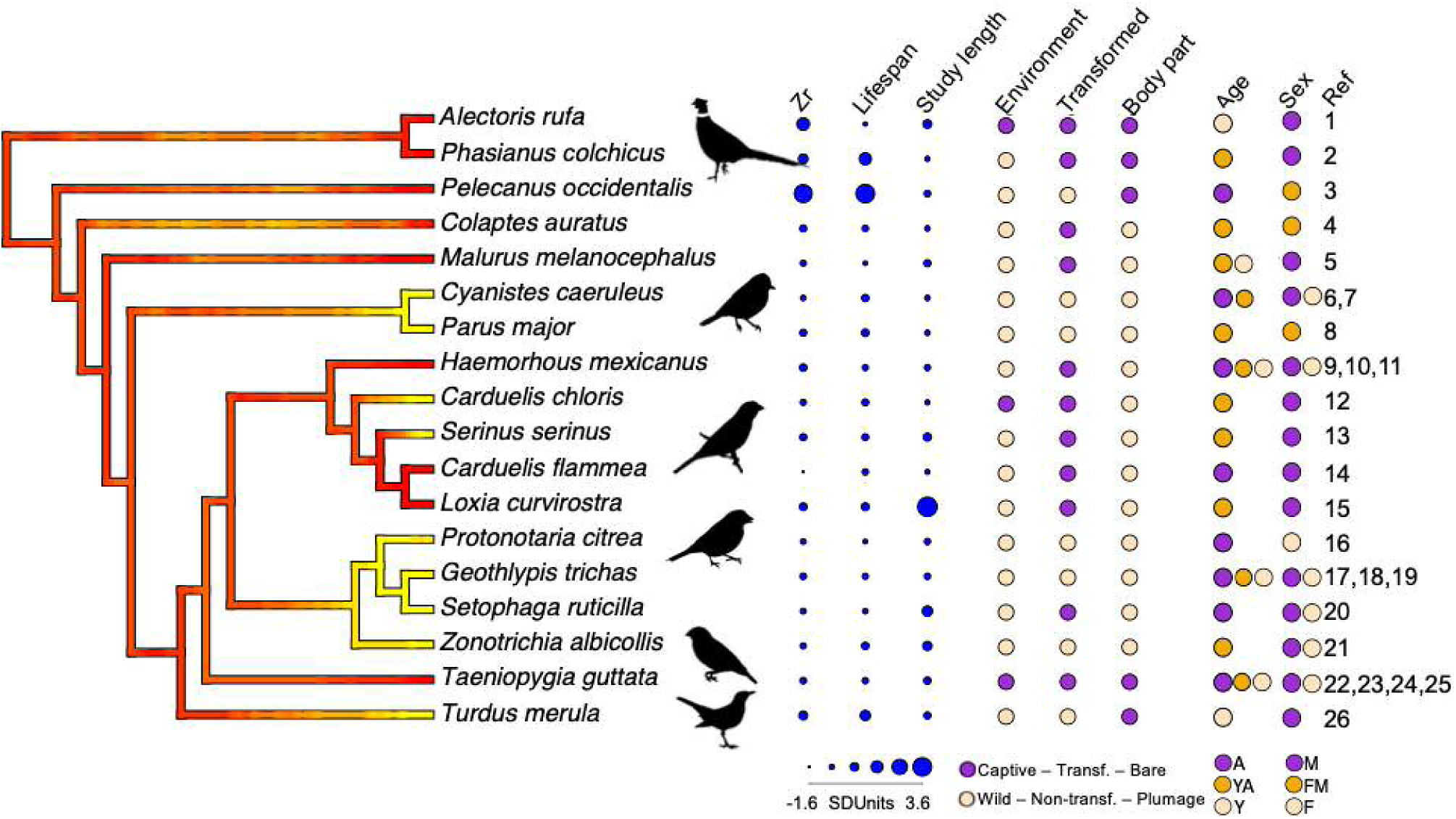
Graphical summary of the phylogenetic distribution of the data in the meta-analysis. The colour of the phylogeny represents the colour of the ornament (red vs. yellow). Continuous variables in blue are standardized across all species with effect size Zr, lifespan measured as the species’ maximum lifespan, and monitoring reflecting the length of the study. Categorical variables with two levels are environment (captivity vs. wild), carotenoid type (transformed vs. dietary) and body (beak vs. plumage). Categorical variables with more levels are sex (male (M), female & male (FM), female (F)) and age at measurement (adult (A), yearling & adult (YA), yearling (Y)). Drawings from www.phylopic.org. References: 1: (Cantarero et al. 2019), 2: (Papeschi and Dessì-Fulgheri 2003), 3: (Jaques 2009); 4: (Wiebe and Bortolotti 2002), 5: (Karubian et al. 2008), 6: (Doutrelant et al. 2008), 7: (Doutrelant et al. 2012), 8: (Hõrak et al. 2001), 9: (Hill 1991), 10: (Hill 1993), 11: (Nolan et al. 1998), 12: (Hõrak and Männiste 2016), 13: (Figuerola and Senar 2007), 14: (Van Oort and Dawson 2005), 15: (Fernández-Eslava et al. 2021), 16: (Slevin et al. 2019), 17: (Dunn et al. 2010), 18: (Freeman-Gallant et al. 2010), 19: (Freeman-Gallant et al. 2014), 20: (Marini et al. 2015), 21: (Grunst et al. 2017), 22: (Price and Burley 1994), 23: (Simons et al. 2012b), 24: (Simons et al. 2016), 25: (Romero-Haro et al. 2025), 26: (Gregoire et al. 2004).

We constructed two phylogenies of the 18 species in this study. First, we implemented the tree from the Open Tree of Life (Hinchliff et al. 2015) with the package ‘rotl’ (Michonneau et al. 2016) and with the branch lengths set following Grafen (1989), by using the function ‘compute.brlen’ from the package ‘ape’ (Paradis and Schliep 2019). Second, we used the recent tree by McTavish et al. 2025 with the package ‘clootl’ (Miller et al. 2025). For both trees, we identified whether the model contained a phylogenetic signal lambda (λ) (Freckleton et al. 2002; Hadfield and Nakagawa 2010), which represents the ratio of the variance explained by phylogeny to the total variance explained by the model. Hence, its value ranges from 0 (no phylogenetic signal) to 1 (all variance explained by phylogeny).

We ran the models using a Bayesian approach with the ‘brm’ function from the ‘brms’ package (Bürkner 2017). In the analyses, we weighted each effect size by the 95% CI of the Zr estimate or by the inverse of the sample size 1/(N-3), where N is the number of individuals, which yielded consistent results. We present the results based on the sample size. We used weakly informative priors and ran four chains with 1,010,000 iterations each, a burn-in of 10,000 and a thinning of 100, always resulting in a posterior effective sample size of more than 15,000. All Rhats were 1.00, Pareto-k-diagnostics, and visual inspection of the trace plots and the potential scale reduction factor showed that simulations had run correctly (Bürkner 2017).

We evaluated the relative fits of the Bayesian model on the data using the leave-one-out (LOO) cross-validation approach (Vehtari et al. 2017) and compared the models’ relative weight with the functions ‘loo’ and ‘model_weights’ of the package ‘brms’ (Bürkner 2017). Briefly, a model’s weight reflects the probability that it will make the best predictions on new data, conditional on the alternative models considered, with the weights of all models summing to 1. We determined the statistical significance of the fixed effects based on the overlap of the coefficients’ 95% CI with 0.

There is no standard approach for the meta-analysis of quadratic associations (three studies; Table S1). Hence, we included all effect sizes, both pre- and post-peak. Note that quadratic associations can emerge from a sign change or when the association between colour and survival approaches zero in part of the colour range. As a sensitivity analysis, we checked the robustness of our conclusions by repeating the analysis using studies that reported only linear effect sizes. We assessed publication bias based on visual inspection of funnel plots of the residuals of the phylogenetic model in Table S2A (Figure S4), the Egger’s test with the function ‘regtest’ of the package ‘metafor’ (Viechtbauer 2010) and using the trim and fill method (Duval and Tweedie 2000) with the function ‘trimfill’ and estimator “R0” of the package ‘metafor’ (Viechtbauer 2010). None of these approaches indicated publication bias (regtest: z = 0.41, p = 0.71; trimfill: z = -0.31, p = 0.75).

## 3. Results

### 3.1 Data composition

We obtained a dataset comprising 26 studies and 57 effect sizes from 18 avian species testing a correlation between carotenoid-based colouration and survival or longevity (Figure 1, Figure 2, Table S1). The majority of the effect sizes (43 out of 57; 75%) came from wild studies, while the rest (14; 25%) was from captive studies (Figure 2, Figure S2, Table S1).

Half of all effect sizes (30 or 53%) were for males, 23 (40%) for females, and 4 (7%) pooled the sexes (Figure 1, Figure 2, Figure S2, Table S1). For six species, we had data on both sexes separately (Figure 2, Table S1, see section 3.4 sensitivity analyses). Most effect sizes (40 or 70%) were based on plumage colouration, and 17 (30%) were based on bare parts colouration (Figure 2, Figure S2, Table S1). Note that the bare parts were mostly beaks, except for the red wattle of the male common pheasants. Most effect sizes (32 or 56%) were based on yellow ornaments, while 25 (44%) were red ornaments (Figure 2, Figure S2, Table S1). Almost half of the effect sizes were based on coloured traits produced by dietary carotenoids (27 or 47%) while the other half were based on ornaments generated by transformed carotenoids (30 or 53%; Figure 2 and S2, Table S1).

Most ornaments were measured in adults (22 or 39%) or combined both yearlings and adults (25 or 44%), while 10 (18%) effect sizes were from yearlings only (Figure 2, Figure S2, Table S1). The average lifespan of all species was 12.5 years (median = 11.6 years), and the average monitoring time of all studies was 4 years (median = 3 years, Figure 2, Table S1).

### 3.2 Phylogenetic variance

We identified whether phylogeny explained a significant proportion of the variation in effect size between carotenoid-based ornament colouration and survival. Models with and without phylogeny fitted the data almost equally well, with respective loo weights of 0.42 and 0.58 (Table S2A vs. S2B, respectively). The phylogeny random term explained a significant proportion of the variance in effect sizes (SD: 0.18, 95% CI: 0.01 – 0.52, Table S2A). This was larger than species (which accounts for the species’ repeated measures; SD: 0.13, 95% CI: 0.09 – 0.34, Table S2A) and as large as the residual variance (SD: 0.16, 95% CI: 0.09 – 0.23, Table S2A). As a result, there was some phylogenetic signal (lambda) at 0.39, albeit with large credibility intervals (95% CI: 0.00 – 0.90, Table S2A). The role of phylogeny was consistent in the full and final models, with the only statistically significant variable sex (see below; SD: 0.15, 95% CI: 0.01 – 0.35; lambda: 0.41, 95% CI: 0.00 – 0.86; loo weights: 0.42 vs 0.42; Table S3A vs. S3B, respectively). Phylogenetic models per predictor variable gave variance estimates of the random effects that were highly consistent across variables and with the full and final models (Tables S2, S3, S4). Hence, we report the results of models that included phylogeny, noting that models with and without phylogeny yielded the same conclusions (full models: Table S2A vs. S2B; final models: Table S3A vs. S3B).

### 3.3 Fixed effects

The mean effect size of our data was 0.15, with the 95% CI overlapping with zero (95% CI: -0.04 – 0.40, Figure 3B, Table S4), indicating a tendency towards a positive association between ornament expression and survival. To understand which variables affected the effect size of the association between colouration and survival, we used two complementary analytic approaches. First, we ran a phylogenetic model with all predictor variables, which revealed one statistically significant variable, sex, for the level males (Table S2A). Second, to obtain the effect size per level, we ran a phylogenetic model for each variable (Figure 3A & B, Table S4A). Both approaches yielded similar results, with effect sizes that were statistically significant and positive for males at 0.20 (95% CI: 0.07 – 0.31) but not for females (Zr = -0.04, 95% CI: -0.23 – 0.10).

**Figure 3.**
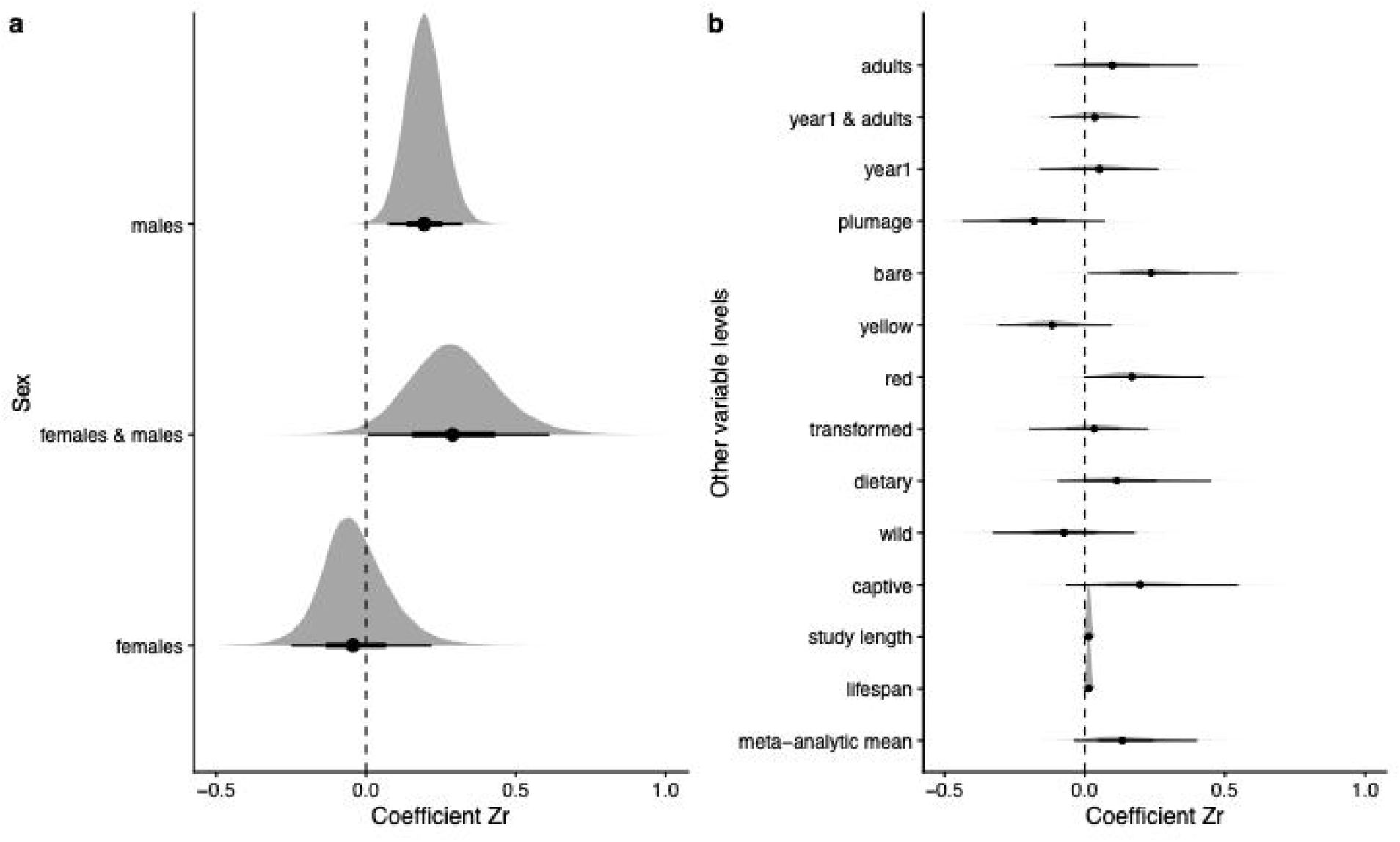
Posterior distributions per variable level of the phylogenetic models analysing the effect sizes between ornament colouration and survival. Grey areas show posterior distributions for (A) the 3 sex levels and (B) all other variable levels. Under the grey distribution, the black dot represents the posterior mean, and the thick and thin horizontal black lines indicate the 66% and 95% credibility intervals, respectively. The vertical dashed line shows an effect size of 0. Level-specific sample sizes and model specifications are shown in the Supplementary Table S4.

To test whether sexual signal characteristics influence the effect size between ornament expression and survival, we included the type of carotenoid in the ornament (dietary untransformed vs. transformed carotenoids, Figure 2), its colour (yellow vs. red, Figure 2) and the body part of the colouration (bare part vs. feathers, Figure 2). In the models per fixed effect (Table S4A), bare parts and red ornaments have 95% credible intervals that might indicate statistically significance, but their significance disappears when (i) models include sex as a covariate (Table S4B & C) and (ii) when excluding the California brown pelicans (*Pelecanus occidentalis*; see sensitivity analyses below). Hence, none of these variables were statistically significant or showed a consistent influence (Figure 3B and S5, Tables S2, S3, S4 and S8).

In addition to the aforementioned fixed effects, the models also included variables accounting for study conditions: study environment (captivity vs. free-living), age at measurement (first-year, adult or first-year and adults combined), species maximum lifespan (obtained from the AnAge database; de Magalhaes and Costa, 2009), or study length (Figure 2). None of these variables were statistically significant (Figure 3B, Tables S2, S3, and S4).

Because the results indicate that carotenoid-based colouration was informative of longevity only for males, we tested for a sex-specific influence of the carotenoid type (dietary vs. transformed), colour (yellow vs. red), or body part (bare part vs. plumage). None of the interactions of these terms with sex were statistically significant (Table S5A and S5B). Hence, there was no evidence that any of these factors significantly affected the sex-specific informational value of the carotenoid-based colouration on survival or longevity.

### 3.4 Sensitivity analyses

We tested the robustness of our conclusions in several ways. First, because there are no specific guidelines for the meta-analysis of quadratic associations, we repeated all analyses with studies that reported linear effect sizes only (49 instead of 57 effect sizes on 17 instead of 18 species, Figure S3), which gave similar results as those described above for both the full model and final model (Table S6A and S6B). Second, in our dataset, there are some species for which information was only available for one sex or where females did not possess a carotenoid-based coloured trait (Figure 2). To understand whether the sex-specific difference in effect sizes arises from these differences, we repeated the final model for the six species containing data on both sexes separately with 29 effect sizes, which gave conclusions consistent with those in Table S2 (females coeff: -0.11, 95% CI: -0.38 – 0.16; males coeff: 0.22, 95% CI: 0.07 – 0.37; Table S7A and S7B), despite it was conducted with a reduced subset of samples. Third, the study on California brown pelicans (*Pelecanus occidentalis*) has a large effect size and longevity, potentially making it an influential data point. Hence, we repeated the full, final and level-specific models excluding pelicans, which again confirmed the sex-specific result (Figure S5, Table S8A, S8B, S8C). These sensitivity analyses also revealed that other variable levels that were (close to) statistically significance in Table S4 became non-significant (Figure S5, Table S8). Fourth, the phylogenetic tree of birds was recently updated by McTavish et al. (2025). Therefore, we repeated the full and final models generated with the traditional tree (Table S2A and S2B) and using the recent tree by McTavish et al. (2025). Analyses with both trees yielded consistent conclusions (Table S9A and S9B). Hence, all sensitivity analyses supported a positive association between carotenoid-based ornament colouration and survival in males but not in females.

## 4. Discussion

Our results revealed a positive relationship between the level of expression of carotenoid-based ornaments and survival among avian species in males but not in females. The finding in males thus supports the reliability of carotenoid-based colourations as signals revealing the bearer’s fitness potential. However, the connection was sex-specific, with more colourful males, but not females, living longer. This sex-specific result contrasts with two meta-analytic studies, also in birds, which found a positive relationship between the expression level of ornaments (including carotenoid-based colourations) and other fitness-related variables in both males and females (Weaver et al. 2018; Nolazco et al. 2022). However, while the former study did not test the association with survival, the latter did not explicitly examine carotenoid-based ornaments but all sexual ornament types together and did not find any correlation with survival in either sex. Our meta-analysis considers a larger number of species with carotenoid-based traits (18 vs. 6) than Nolazco et al. (2022) and is the first to test specifically the relationship between carotenoid-based ornament expression and survival or longevity. Our results indicate the association is common among avian species and is sex-specific.

The role of carotenoid-based colouration in signalling longevity has been previously suggested from results in fish and birds (Pike et al. 2007; Freeman-Gallant et al. 2011; Romero-Haro et al. 2025). The positive association between carotenoid-based colouration and longevity could arise in various ways. It could be explained by the role of carotenoids in immune function (Chew and Park 2004; Milani et al. 2017) and antioxidant defences (Svensson and Wong 2011; Nabi et al. 2020; but see Koch et al. 2018). Indeed, oxidative stress (i.e., the imbalance between antioxidant levels and production of reactive oxygen species by cell metabolism; e.g., Harman 2006; Speakman et al. 2015) is an important trigger of ageing-related diseases and, consequently, reduced longevity (Barja 2002; Harman 2006; Jiménez 2018; Luo et al. 2020). If carotenoids are a limited resource (Endler 1980; Kodric-Brown 1985; Hill 1990; but see e.g. Hadfield and Owens 2006 and Simons et al. 2014) and given their role in homeostasis-related functions (e.g. Nabi et al. 2020), our findings would support the hypothesis that the reliability of carotenoid-based signals is maintained by a trade-off between allocating carotenoids to colouration vs self-maintenance (von Schantz et al. 1999; Møller et al. 2000; Alonso-Alvarez et al. 2008; but see Koch and Hill 2018). We acknowledge, though, that our approach does not allow us to identify the specific functional association between ornament colouration and survival, which remains challenging.

According to the "shared-pathway hypothesis" (Hill 2011; Hill et al. 2019), we predicted that colourations based on transformed carotenoids would be more strongly correlated with survival than those based on untransformed dietary pigments. In the analysis that included all the species, we did not detect significant differences in effect sizes between these traits. However, when removing the only seabird species (i.e., Californian brown pelican; Jaques et al. 2019) from the analysis, the effect sizes of red ornaments or those coloured by transformed carotenoids were, on average, larger than those of yellow traits or those produced by untransformed yellow carotenoids, respectively (Figure S2A,B). Seabirds can acquire ketocarotenoids directly from their diet (e.g. arthropods or fish containing high amounts of these red pigments; McGraw 2006). Therefore, its exclusion allowed us to compare better the effect of ornaments coloured by transformed and non-transformed pigments. Hence, considering the weak trend detected when removing the seabird study, we may still speculate that, in the future, a meta-analysis with a larger number of studies could support the prediction. Interestingly, in the meta-analysis by Weaver et al. (2018), the colour intensity due to transformed carotenoids, but not to dietary yellow pigments, was positively correlated with condition- and fecundity-related parameters. That meta-analysis included a similar number of avian species (i.e., 19) as the current study but was restricted to feather colouration in passerine species. In contrast, we examined a broader range of taxa (Figure 2), encompassing both bare parts and plumage colouration (Figure 2). Bare parts based on carotenoid colouration differ from plumage colouration in that they are more dynamic, require sustained production periods, and are usually located farther from the carotenoid-transformation tissues than feathers (Alonso-Alvarez et al. 2022). Therefore, the discrepancies between our study and Weaver et al. (2018) meta-analysis may stem from differences in ornament characteristics, taxa, or, ultimately, the fitness parameters tested.

Female ornaments did not exhibit a significant association with survival (Figure 3). This could be attributed to weaker sexual selection pressure on female carotenoid-based signals (e.g., Dunn et al. 2015; Clark and Rankin 2020). Although some studies support direct sexual or social selection on female ornaments (e.g., Dale et al. 2015; Hare and Simmons 2019; Doutrelant et al. 2020), our findings rather support the view that the higher reproductive investment of females (due to anisogamy) and the derived costs may constrain the evolution of carotenoid-based traits as reliable signals of individual quality in this sex (Fitzpatrick et al. 1995; Morales et al. 2009). A non-excluding possibility is that males likely benefit more than females from signalling, as they presumably maximise their fitness by increasing their mating rates (i.e., Bateman, 1948; see also Collet et al. 2014). In any event, females could gain more benefits than males by choosing mates based on carotenoid-based ornaments, particularly in terms of offspring viability.

Additionally, we suggest that female ornaments could better signal fecundity-associated parameters than longevity. Two meta-analyses in birds, the most studied taxon in this context, found a positive correlation between the expression level of ornaments and several fecundity-related parameters that was consistent in both sexes (Weaver et al. 2018; Nolazco et al. 2022). Another meta-analysis by Hernández et al. (2021) performed exclusively in females also showed a significant positive association between carotenoid-based colourations and clutch size. In contrast, regarding longevity, only Nolazco et al. (2022) conducted a meta-analysis testing the connection between avian ornaments and survival, with no significant association in either sex. Nolazco et al. (2020) excluded species without sexual dichromatism and involved a broader range of ornaments, including melanin-based, structural colourations, achromatic and morphological traits. However, their data consisted of fewer (only six) species where carotenoid-based ornaments were measured and correlated with survival (Nolazco et al. 2022). Therefore, our dataset allowed us to specifically test the link between carotenoid-based traits and survival with greater power, which may explain differences between the two studies.

Our sex-specific conclusion is consistent with a recent meta-analysis of meta-analyses that tested the relationship between survival and the conspicuousness of a wide range of putative ornaments across various taxa, including vertebrates and invertebrates, and is not specific to carotenoid-based sexual signals (i.e., Pollo et al. 2025). Moreover, we extracted data directly from the primary literature, resulting in a larger sample for testing the carotenoid-based ornament level x survival correlation than that of Pollo et al. (2025), which relied largely on data from Nolazco et al. (2022). Furthermore, our focus on traits produced by carotenoid pigments allowed us to explore the mechanism-based hypothesis proposed by Hill et al. (2019). Our results suggest that sex-specific associations between carotenoid-based ornaments and survival can be generalized across ornament production mechanisms.

Sexual dimorphism has traditionally been interpreted as the result of stronger sexual selection on male traits and natural selection on female traits, undervaluing the influence of direct sexual selection on females (e.g., Amundsen 2000; Price 2015; Doutrelant et al. 2020). Here, among the 18 species with available data, the carotenoid-based ornament tested in the analyses was absent in females only in two species: *Phasianus colchicus* and *Carduelis flammea*. The remaining species showed moderate sexual differences in ornament expression, suggesting that the coloured traits may have evolved through mutual sexual (or social) selection in these cases (Johnstone 1996; Kokko and Johnstone 2002). However, experimental demonstrations of mate choice based on carotenoid-based colourations in these species are rare and primarily focused on female preferences (i.e., Burley and Coopersmith 1987; Karubian 2002; Dunn et al. 2010; Alonso-Alvarez et al. 2012; Toomey and McGraw 2012; Trigo et al. 2024). Currently, only six species have available effect sizes for both sexes (Table S1), and this subsample yielded results consistent with those of the full dataset (Table S7A and S7B). Therefore, the conclusions are robust to species-related variation in sex-specific signals.

Finally, the lack of association between carotenoid-based ornament level and female survival could also be due to the shorter longevity of females compared to males in avian species (Xirocostas et al. 2020). In a correlation, a small range of variation in one variable can make it more difficult to detect a significant association (e.g., Bland and Altman 2011). In that case, the sex-related differences in the correlation between carotenoid-based colouration and survival would be a consequence of sex-related differences in life history.

To conclude, the relevance of our results lies in the sex-specific association between carotenoid-based signalling and longevity, for which we provide several non-exclusive explanations: (i) weaker sexual selection in females likely due to higher reproductive investment-based constraints than males, (ii) female ornaments signalling fecundity rather than longevity, and (iii) sex-specific life history variation. Our results do not provide support for the hypothesis that ornaments coloured by enzymatically transformed carotenoids are more reliable signals of longevity than those produced by dietary pigments. Instead, they suggest that avian carotenoid-based ornaments are reliable signals of longevity in males, independent of the proximate production mechanism.

## Acknowledgements

This work was supported by projects PID2019-109303GB-I00 to (C.A.A.) and PID2022-139166NB-I00 (to J.M.) both funded by Ministerio de Ciencia e Innovación of the Spanish government (MCIN/AEI/ 10.13039/501100011033 program), and the latter also by ‘ERDF A way of making Europe’ program). A.A.R.-H. was supported by a CAM Talent Attraction grant (2024-T1/ECO-31271). We also thank the funding from the Turku Collegium for Science, Medicine and Technology.

## Colour measures and assignment to carotenoid type

The expression level of the carotenoid-based ornaments was determined from trait size or colour intensity measures. Variables included subjective colour rankings obtained by the researchers or objective measures from digital cameras or colourimeters. The trait brightness was excluded from our analyses because it is strongly influenced by tissue’s physical microstructures, which can be altered by abrasion and wear, making it difficult to interpret in terms of carotenoid content (e.g. Weaver et al. 2018). The coloured ornament analysed in each species was assigned to one of two categories: (1) colourations produced by transformed carotenoids, including both red ketocarotenoids and yellow canary xanthophylls and (2) ornaments coloured only by non-transformed dietary carotenoids (e.g., lutein, zeaxanthin, b-carotene; e.g., McGraw 2006; Weaver et al. 2018). The carotenoid type of the ornaments of 15 species in Table S1 was determined from studies that included chromatographic measurements (i.e., Faivre et al. 2003; McGraw 2006 and references therein; Thomas et al. 2014; García-de Blas et al. 2016; Khalil et al. 2023). The remaining species were assigned following the most parsimonious explanation. Regarding the yellow plumage of prothonotary warblers (*Protonotaria citrea*), this species is phylogenetically related to *Vermivora* (Lovette et al. 2010), where yellow plumages are only produced by dietary lutein (data in McGraw 2006). Male white-throated sparrows (*Zonotrichia albicollis*) show yellow plumage patches on their heads, which are attributed to carotenoids (Grunst et al. 2017). We conservatively assigned its colouration to dietary untransformed carotenoids, consistent with the aforementioned literature and the most common option among passerines (McGraw 2006). We excluded a study (i.e., Burke et al. 2022) on bobolinks (*Dolichonyx oryzivorus*), a passerine species with a yellow plumage patch in the nape of males only, since we could not find any published study reporting analyses on its biochemical composition, although unpublished results seem to support the idea that it is created by pheomelanins (Kevin J McGraw, personal communication).

**Figure S1.**
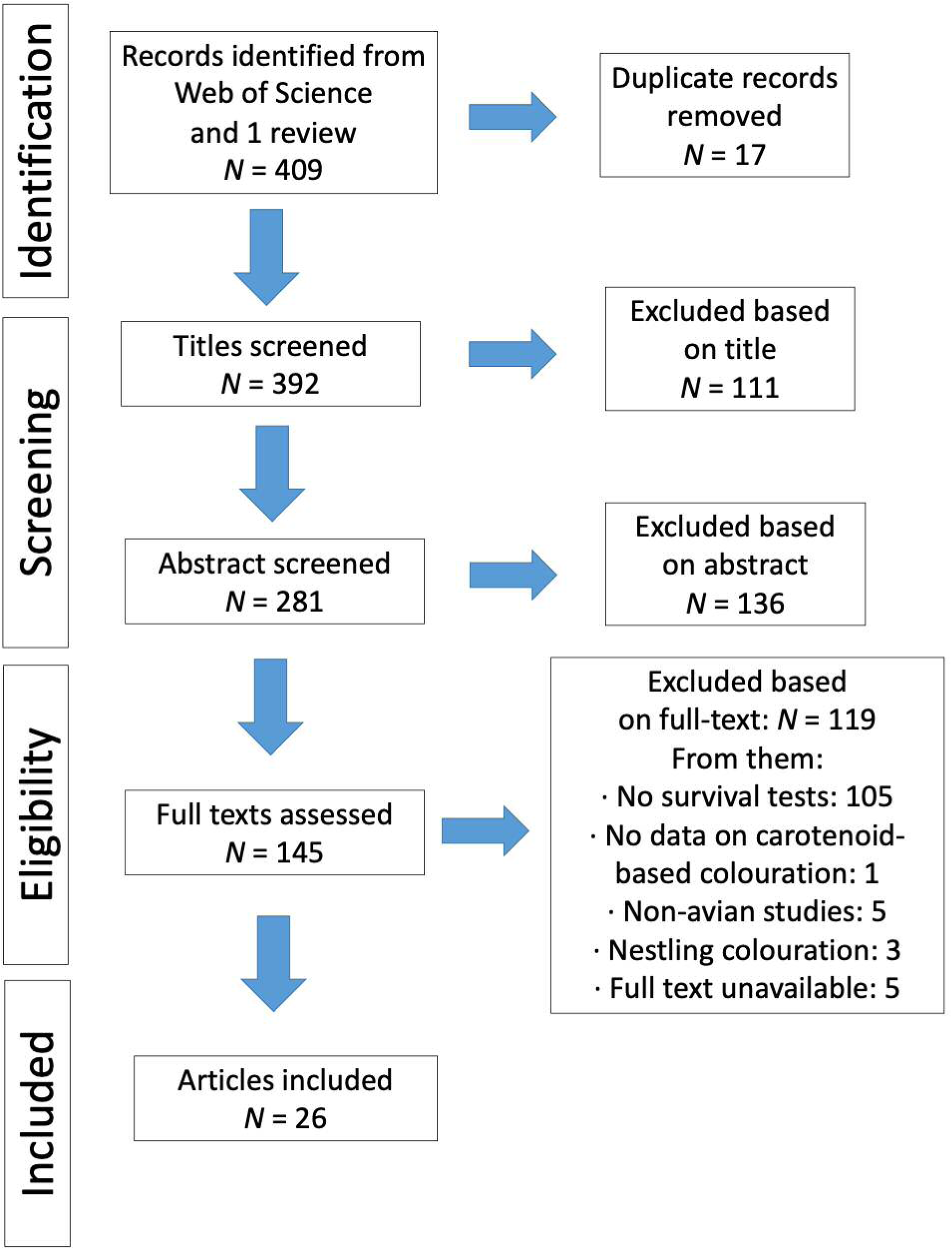
Prisma flow chart, following the PRISMA guidelines (Page et al. 2021), of the literature search for associations between sexual colouration and survival in birds.

**Figure S2.**
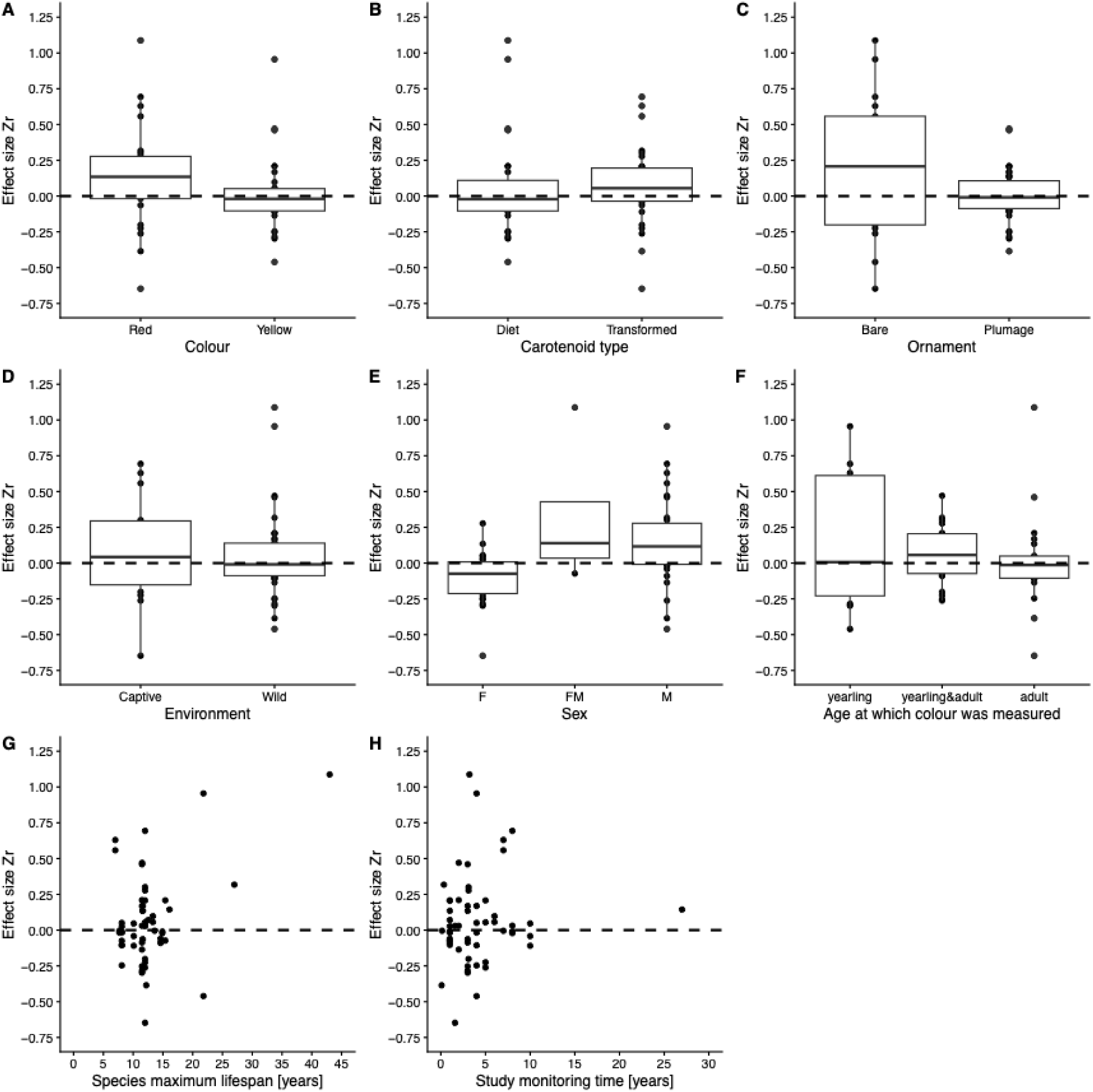
Distribution of the 57 effect sizes per covariate. Abbreviations: M = males, F = females, FM = females and males combined.

**Figure S3.**
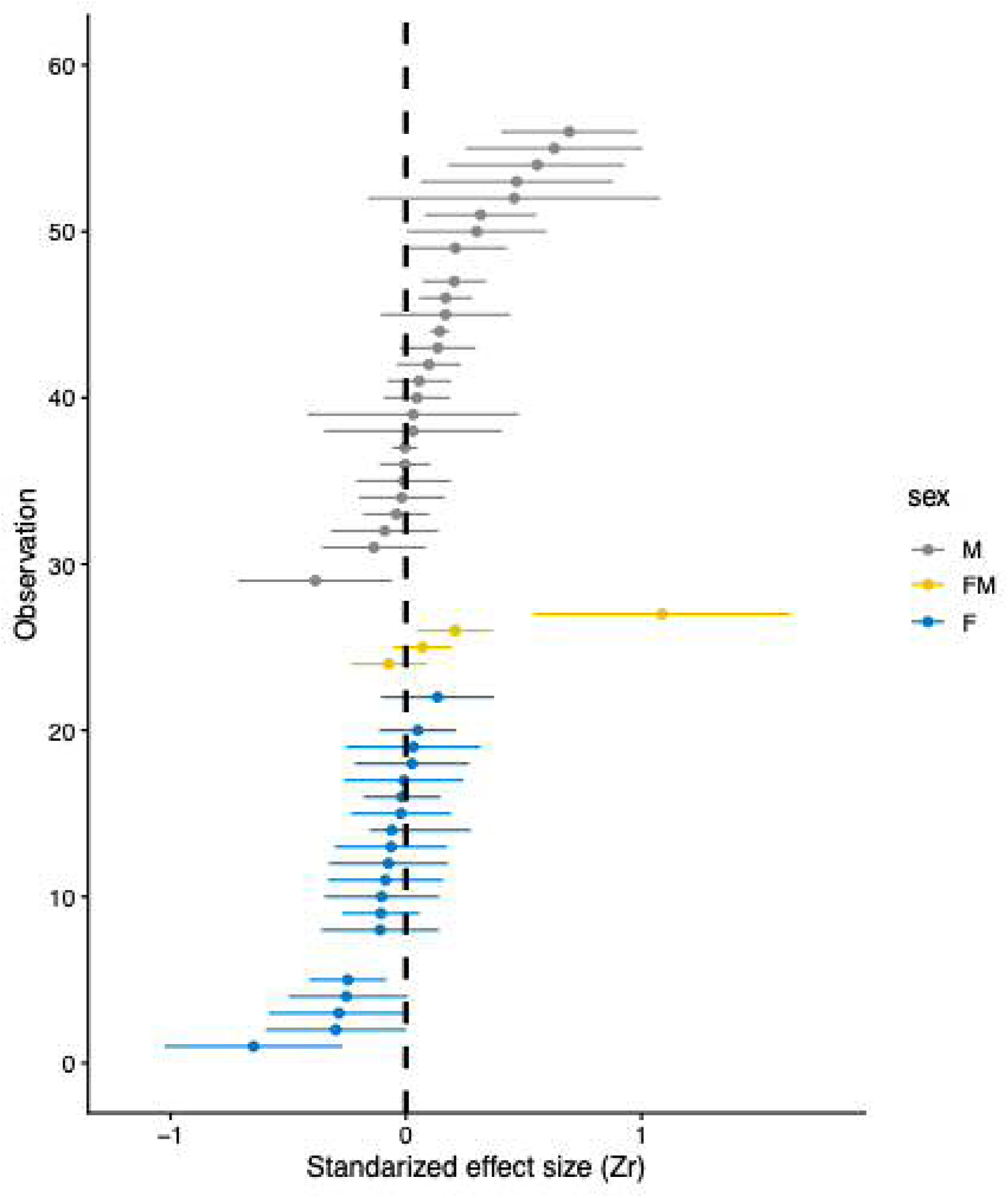
Distribution of the 49 linear effect sizes per sex group. Effect sizes are ranked in the same order as for Figure 1. Abbreviations: M = males, F = females, FM = females and males combined.

**Figure S4.**
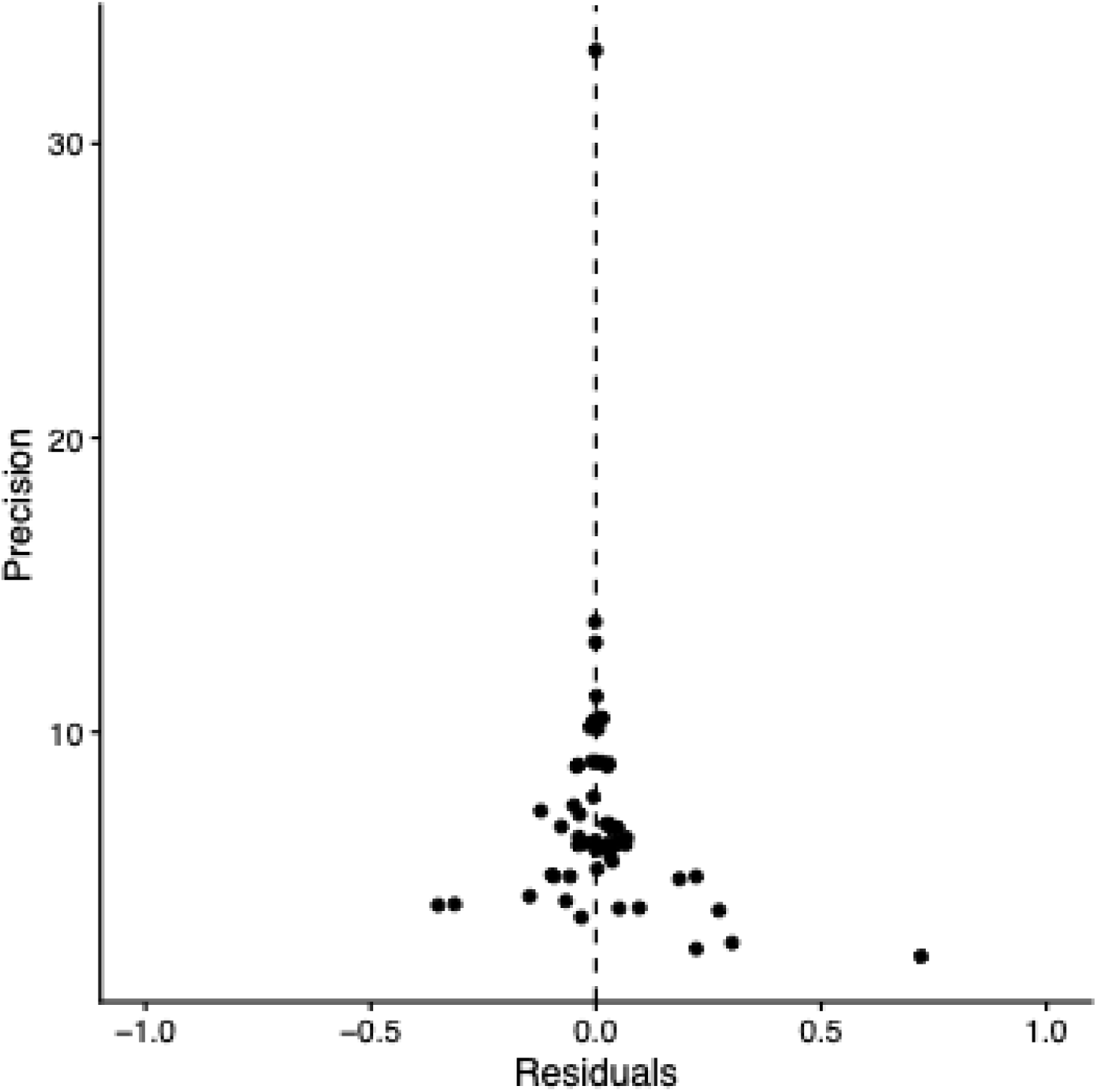
Funnel plot showing the residuals of the full phylogenetic model in Table S2A.

**Figure S5.**
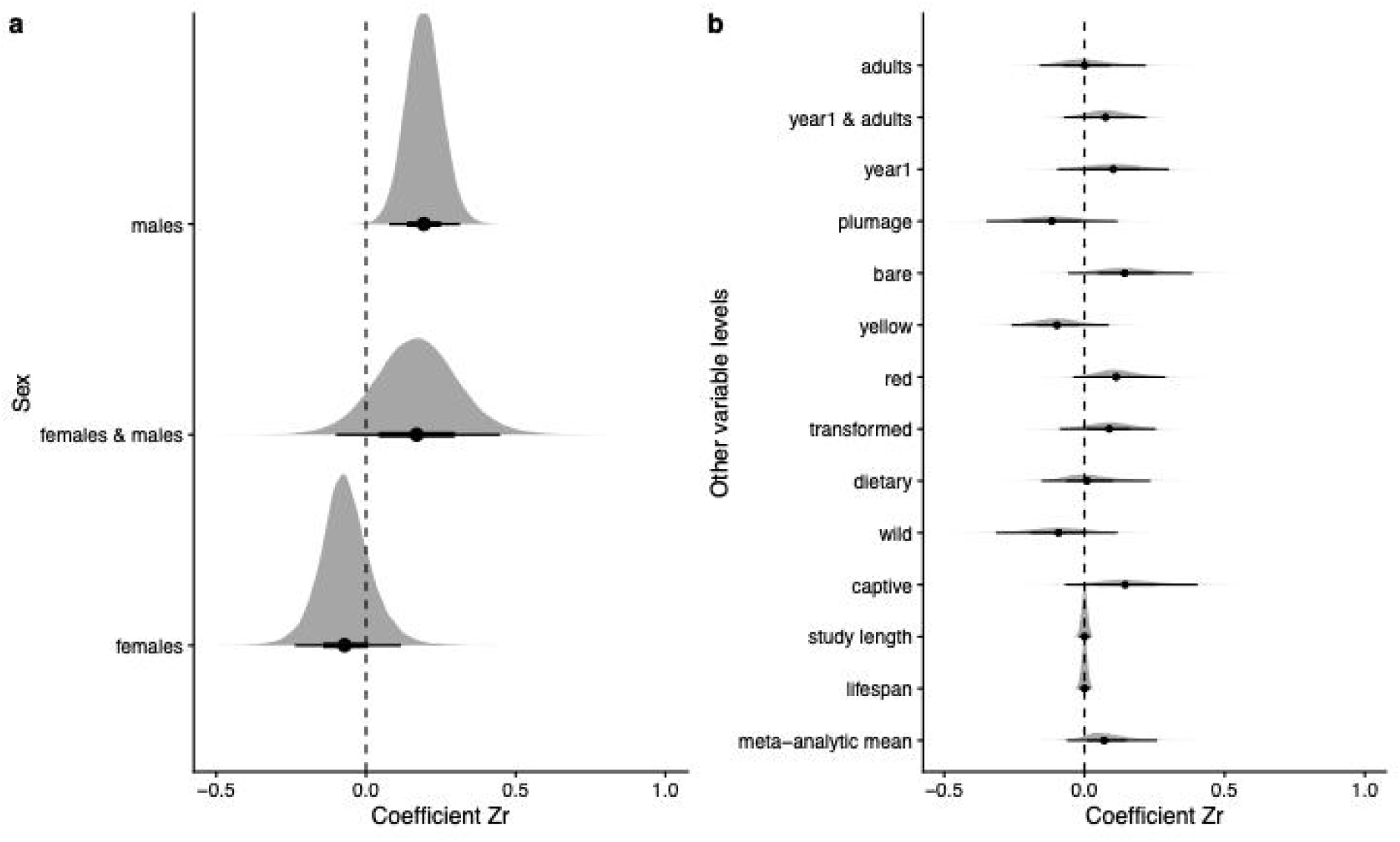
Posterior distributions per variable level of models as in Figure 3, but using a dataset without the California brown pelicans (*Pelecanus occidentalis*).

**Table S1.**
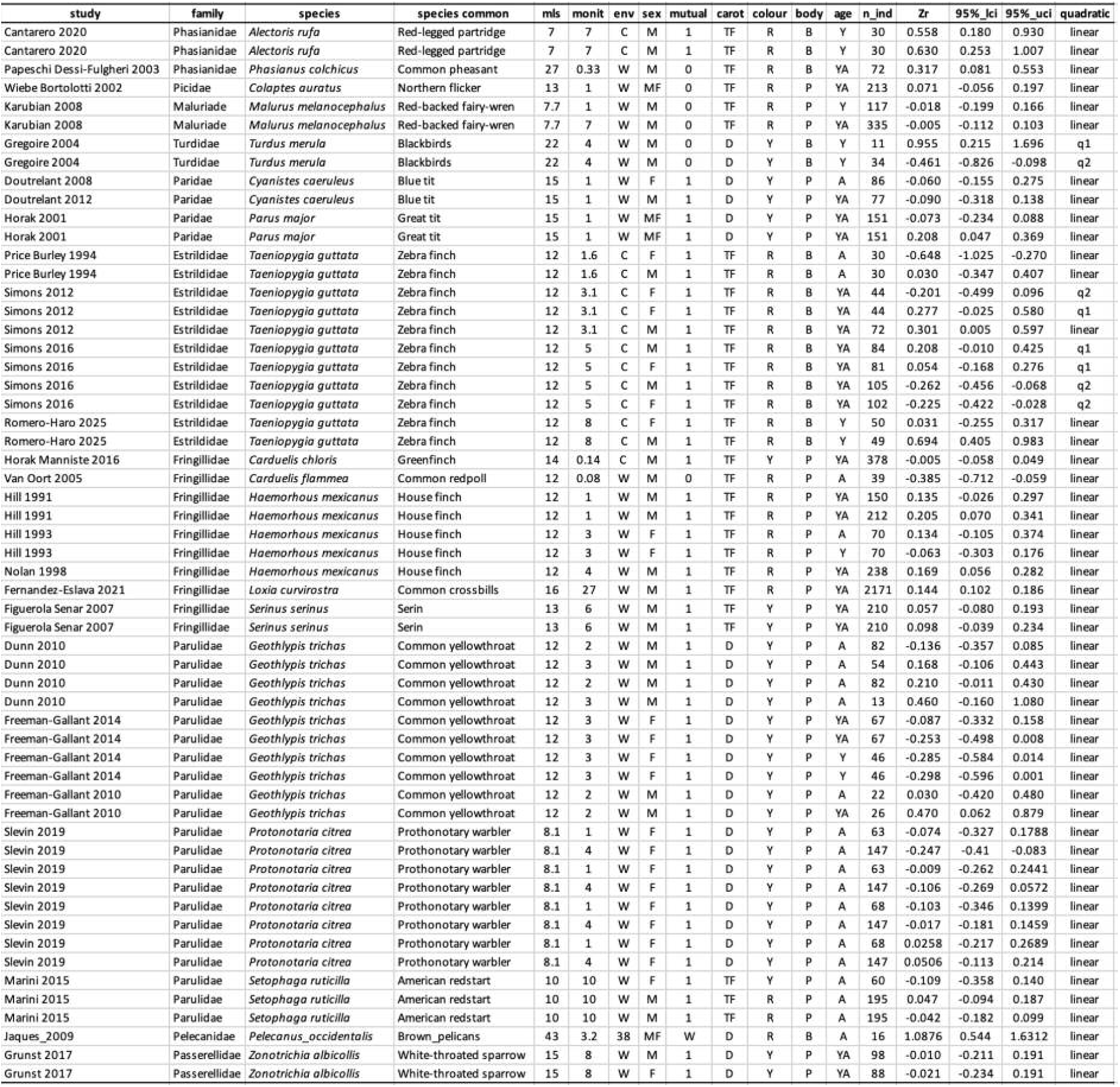
Data extracted from the 26 studies following the literature search in Fig. S1. Abbreviations: mls = maximum lifespan; monit = study length in years; env = environment, c = captive, w = wild; sex: m = males, mf = males and females, f = females; mutual: whether both sexes display ornaments (1) or males only (0); carot = carotenoid, tf = transformed, d = dietary; colour: r = red, y = yellow; body: b = bare parts, p = plumage; age: a = adult, ya = yearling and adult, y = yearling, nest = nestling; n_ind = number of individuals; Zr = effect size; 95%_lci = lower 95% confidence interval of Zr; 95%_uci = upper 95% confidence interval of Zr; quadratic = whether the effect size was part of a quadratic association, q1 = pre-maximum effect size, q2 = post-maximum effect size.

| study | family | species | species common | mls | monit | env | sex | mutual | carot | colour | body | age | n_ind | Zr | 95%_lci | 95%_uci | quadratic |
| --- | --- | --- | --- | --- | --- | --- | --- | --- | --- | --- | --- | --- | --- | --- | --- | --- | --- |
| Cantarero 2020 | Phasianidae | *Alectoris rufa* | Red-legged partridge | 7 | 7 | C | M | 1 | TF | R | B | Y | 30 | 0.558 | 0.180 | 0.930 | linear |
| Cantarero 2020 | Phasianidae | *Alectoris rufa* | Red-legged partridge | 7 | 7 | C | M | 1 | TF | R | B | Y | 30 | 0.630 | 0.253 | 1.007 | linear |
| Papeschi Dessi-Fulgheri 2003 | Phasianidae | *Phasianus colchicus* | Common pheasant | 27 | 0.33 | W | M | 0 | TF | R | B | YA | 72 | 0.317 | 0.081 | 0.553 | linear |
| Wiebe Bortolotti 2002 | Picidae | *Colaptes auratus* | Northern flicker | 13 | 1 | W | MF | 0 | TF | R | P | YA | 213 | 0.071 | -0.056 | 0.197 | linear |
| Karubian 2008 | Maluriade | *Malurus melanocephalus* | Red-backed fairy-wren | 7.7 | 1 | W | M | 0 | TF | R | P | Y | 117 | -0.018 | -0.199 | 0.166 | linear |
| Karubian 2008 | Maluriade | *Malurus melanocephalus* | Red-backed fairy-wren | 7.7 | 7 | W | M | 0 | TF | R | P | YA | 335 | -0.005 | -0.112 | 0.103 | linear |
| Gregoire 2004 | Turdidae | *Turdus merula* | Blackbirds | 22 | 4 | W | M | 0 | D | Y | B | Y | 11 | 0.955 | 0.215 | 1.696 | q1 |
| Gregoire 2004 | Turdidae | *Turdus merula* | Blackbirds | 22 | 4 | W | M | 0 | D | Y | B | Y | 34 | -0.461 | -0.826 | -0.098 | q2 |
| Doutrelant 2008 | Paridae | *Cyanistes caeruleus* | Blue tit | 15 | 1 | W | F | 1 | D | Y | P | A | 86 | -0.060 | -0.155 | 0.275 | linear |
| Doutrelant 2012 | Paridae | *Cyanistes caeruleus* | Blue tit | 15 | 1 | W | M | 1 | D | Y | P | YA | 77 | -0.090 | -0.318 | 0.138 | linear |
| Horak 2001 | Paridae | *Parus major* | Great tit | 15 | 1 | W | MF | 1 | D | Y | P | YA | 151 | -0.073 | -0.234 | 0.088 | linear |
| Horak 2001 | Paridae | *Parus major* | Great tit | 15 | 1 | W | MF | 1 | D | Y | P | YA | 151 | 0.208 | 0.047 | 0.369 | linear |
| Price Burley 1994 | Estrildidae | *Taeniopygia guttata* | Zebra finch | 12 | 1.6 | C | F | 1 | TF | R | B | A | 30 | -0.648 | -1.025 | -0.270 | linear |
| Price Burley 1994 | Estrildidae | *Taeniopygia guttata* | Zebra finch | 12 | 1.6 | C | M | 1 | TF | R | B | A | 30 | 0.030 | -0.347 | 0.407 | linear |
| Simons 2012 | Estrildidae | *Taeniopygia guttata* | Zebra finch | 12 | 3.1 | C | F | 1 | TF | R | B | YA | 44 | -0.201 | -0.499 | 0.096 | q2 |
| Simons 2012 | Estrildidae | *Taeniopygia guttata* | Zebra finch | 12 | 3.1 | C | F | 1 | TF | R | B | YA | 44 | 0.277 | -0.025 | 0.580 | q1 |
| Simons 2012 | Estrildidae | *Taeniopygia guttata* | Zebra finch | 12 | 3.1 | C | M | 1 | TF | R | B | YA | 72 | 0.301 | 0.005 | 0.597 | linear |
| Simons 2016 | Estrildidae | *Taeniopygia guttata* | Zebra finch | 12 | 5 | C | M | 1 | TF | R | B | YA | 84 | 0.208 | -0.010 | 0.425 | q1 |
| Simons 2016 | Estrildidae | *Taeniopygia guttata* | Zebra finch | 12 | 5 | C | F | 1 | TF | R | B | YA | 81 | 0.054 | -0.168 | 0.276 | q1 |
| Simons 2016 | Estrildidae | *Taeniopygia guttata* | Zebra finch | 12 | 5 | C | M | 1 | TF | R | B | YA | 105 | -0.262 | -0.456 | -0.068 | q2 |
| Simons 2016 | Estrildidae | *Taeniopygia guttata* | Zebra finch | 12 | 5 | C | F | 1 | TF | R | B | YA | 102 | -0.225 | -0.422 | -0.028 | q2 |
| Romero-Haro 2025 | Estrildidae | *Taeniopygia guttata* | Zebra finch | 12 | 8 | C | F | 1 | TF | R | B | Y | 50 | 0.031 | -0.255 | 0.317 | linear |
| Romero-Haro 2025 | Estrildidae | *Taeniopygia guttata* | Zebra finch | 12 | 8 | C | M | 1 | TF | R | B | Y | 49 | 0.694 | 0.405 | 0.983 | linear |
| Horak Manniste 2016 | Fringillidae | *Carduelis chloris* | Greenfinch | 14 | 0.14 | C | M | 1 | TF | Y | P | YA | 378 | -0.005 | -0.058 | 0.049 | linear |
| Van Oort 2005 | Fringillidae | *Carduelis flammea* | Common redpoll | 12 | 0.08 | W | M | 0 | TF | R | P | A | 39 | -0.385 | -0.712 | -0.059 | linear |
| Hill 1991 | Fringillidae | *Haemorrhous mexicanus* | House finch | 12 | 1 | W | M | 1 | TF | R | P | YA | 150 | 0.135 | -0.026 | 0.297 | linear |
| Hill 1991 | Fringillidae | *Haemorrhous mexicanus* | House finch | 12 | 1 | W | M | 1 | TF | R | P | YA | 212 | 0.205 | 0.070 | 0.341 | linear |
| Hill 1993 | Fringillidae | *Haemorrhous mexicanus* | House finch | 12 | 3 | W | F | 1 | TF | R | P | A | 70 | 0.134 | -0.105 | 0.374 | linear |
| Hill 1993 | Fringillidae | *Haemorrhous mexicanus* | House finch | 12 | 3 | W | F | 1 | TF | R | P | Y | 70 | -0.063 | -0.303 | 0.176 | linear |
| Nolan 1998 | Fringillidae | *Haemorrhous mexicanus* | House finch | 12 | 4 | W | M | 1 | TF | R | P | YA | 238 | 0.169 | 0.056 | 0.282 | linear |
| Fernandez-Eslava 2021 | Fringillidae | *Loxia curvirostra* | Common crossbills | 16 | 27 | W | M | 1 | TF | R | P | YA | 2171 | 0.144 | 0.102 | 0.186 | linear |
| Figuerola Senar 2007 | Fringillidae | *Serinus serinus* | Serin | 13 | 6 | W | M | 1 | TF | Y | P | YA | 210 | 0.057 | -0.080 | 0.193 | linear |
| Figuerola Senar 2007 | Fringillidae | *Serinus serinus* | Serin | 13 | 6 | W | M | 1 | TF | Y | P | YA | 210 | 0.098 | -0.039 | 0.234 | linear |
| Dunn 2010 | Parulidae | *Geothlypis trichas* | Common yellowthroat | 12 | 2 | W | M | 1 | D | Y | P | A | 82 | -0.136 | -0.357 | 0.085 | linear |
| Dunn 2010 | Parulidae | *Geothlypis trichas* | Common yellowthroat | 12 | 3 | W | M | 1 | D | Y | P | A | 54 | 0.168 | -0.106 | 0.443 | linear |
| Dunn 2010 | Parulidae | *Geothlypis trichas* | Common yellowthroat | 12 | 2 | W | M | 1 | D | Y | P | A | 82 | 0.210 | -0.011 | 0.430 | linear |
| Dunn 2010 | Parulidae | *Geothlypis trichas* | Common yellowthroat | 12 | 3 | W | M | 1 | D | Y | P | A | 13 | 0.460 | -0.160 | 1.080 | linear |
| Freeman-Gallant 2014 | Parulidae | *Geothlypis trichas* | Common yellowthroat | 12 | 3 | W | F | 1 | D | Y | P | YA | 67 | -0.087 | -0.332 | 0.158 | linear |
| Freeman-Gallant 2014 | Parulidae | *Geothlypis trichas* | Common yellowthroat | 12 | 3 | W | F | 1 | D | Y | P | YA | 67 | -0.253 | -0.498 | 0.008 | linear |
| Freeman-Gallant 2014 | Parulidae | *Geothlypis trichas* | Common yellowthroat | 12 | 3 | W | F | 1 | D | Y | P | Y | 46 | -0.285 | -0.584 | 0.014 | linear |
| Freeman-Gallant 2014 | Parulidae | *Geothlypis trichas* | Common yellowthroat | 12 | 3 | W | F | 1 | D | Y | P | Y | 46 | -0.298 | -0.596 | 0.001 | linear |
| Freeman-Gallant 2010 | Parulidae | *Geothlypis trichas* | Common yellowthroat | 12 | 2 | W | M | 1 | D | Y | P | A | 22 | 0.030 | -0.420 | 0.480 | linear |
| Freeman-Gallant 2010 | Parulidae | *Geothlypis trichas* | Common yellowthroat | 12 | 2 | W | M | 1 | D | Y | P | YA | 26 | 0.470 | 0.062 | 0.879 | linear |
| Slevin 2019 | Parulidae | *Protonotaria citrea* | Prothonotary warbler | 8.1 | 1 | W | F | 1 | D | Y | P | A | 63 | -0.074 | -0.327 | 0.1788 | linear |
| Slevin 2019 | Parulidae | *Protonotaria citrea* | Prothonotary warbler | 8.1 | 4 | W | F | 1 | D | Y | P | A | 147 | -0.247 | -0.41 | -0.083 | linear |
| Slevin 2019 | Parulidae | *Protonotaria citrea* | Prothonotary warbler | 8.1 | 1 | W | F | 1 | D | Y | P | A | 63 | -0.009 | -0.262 | 0.2441 | linear |
| Slevin 2019 | Parulidae | *Protonotaria citrea* | Prothonotary warbler | 8.1 | 4 | W | F | 1 | D | Y | P | A | 147 | -0.106 | -0.269 | 0.0572 | linear |
| Slevin 2019 | Parulidae | *Protonotaria citrea* | Prothonotary warbler | 8.1 | 1 | W | F | 1 | D | Y | P | A | 68 | -0.103 | -0.346 | 0.1399 | linear |
| Slevin 2019 | Parulidae | *Protonotaria citrea* | Prothonotary warbler | 8.1 | 4 | W | F | 1 | D | Y | P | A | 147 | -0.017 | -0.181 | 0.1459 | linear |
| Slevin 2019 | Parulidae | *Protonotaria citrea* | Prothonotary warbler | 8.1 | 1 | W | F | 1 | D | Y | P | A | 68 | 0.0258 | -0.217 | 0.2689 | linear |
| Slevin 2019 | Parulidae | *Protonotaria citrea* | Prothonotary warbler | 8.1 | 4 | W | F | 1 | D | Y | P | A | 147 | 0.0506 | -0.113 | 0.214 | linear |
| Marini 2015 | Parulidae | *Setophaga ruticilla* | American redstart | 10 | 10 | W | F | 1 | TF | Y | P | A | 60 | -0.109 | -0.358 | 0.140 | linear |
| Marini 2015 | Parulidae | *Setophaga ruticilla* | American redstart | 10 | 10 | W | M | 1 | TF | R | P | A | 195 | 0.047 | -0.094 | 0.187 | linear |
| Marini 2015 | Parulidae | *Setophaga ruticilla* | American redstart | 10 | 10 | W | M | 1 | TF | R | P | A | 195 | -0.042 | -0.182 | 0.099 | linear |
| Jaques 2009 | Pelecanidae | *Pelecanus occidentalis* | Brown pelicans | 43 | 3.2 | 38 | MF | W | D | R | B | A | 16 | 1.0876 | 0.544 | 1.6312 | linear |
| Grunst 2017 | Passerellidae | *Zonotrichia albicollis* | White-throated sparrow | 15 | 8 | W | M | 1 | D | Y | P | YA | 98 | -0.010 | -0.211 | 0.191 | linear |
| Grunst 2017 | Passerellidae | *Zonotrichia albicollis* | White-throated sparrow | 15 | 8 | W | F | 1 | D | Y | P | YA | 88 | -0.021 | -0.234 | 0.191 | linear |

**Table S2.** Full models (A) with and (B) without phylogeny. The levels in italics are statistically significant. Abbreviations: SD: Standard deviation explained by the random intercept; Est.Error: standard error of the estimate or SD; l-95%CI: lower 95% credibility interval; u-95%CI: upper 95% credibility interval; Rhat: convergence diagnostic with a value of 1 indicating that chains have mixed properly ESS: effective sample size of the estimate or SD with Bulk_ESS showing the effective sample size for the mean of the distribution posterior distribution and Tail_ESS showing the minimum effective sample sizes for 5% and 95% quantiles of the posterior distribution. Further explanations of the diagnostic Markov Chains in brms are given in Bürkner et al. (2017) and Vehtari et al. (2021).

| **(A) Full model with phylogeny** |  |  |  |  |  |  |  |
| --- | --- | --- | --- | --- | --- | --- | --- |
| **Random terms** | **SD** | **Est.Error** | **l-95%CI** | **u-95%CI** | **Rhat** | **Bulk_ESS** | **Tail_ESS** |
| ~species | 0.13 | 0.09 | 0.01 | 0.34 | 1.00 | 38904 | 39184 |
| ~residual | 0.16 | 0.04 | 0.09 | 0.23 | 1.00 | 39829 | 38974 |
| ~phylogeny | 0.18 | 0.14 | 0.01 | 0.52 | 1.00 | 39941 | 39828 |
| phylogenetic signal lambda | 0.39 | 0.26 | 0.00 | 0.90 |  |  |  |
| **Fixed effects** | **Estimate** | **Est.Error** | **l-95%CI** | **u-95%CI** | **Rhat** | **Bulk_ESS** | **Tail_ESS** |
| Intercept | 0.06 | 0.34 | -0.62 | 0.76 | 1.00 | 39723 | 38306 |
| maximum lifespan | 0.01 | 0.01 | -0.02 | 0.04 | 1.00 | 40273 | 39680 |
| monitoring years | 0.01 | 0.01 | -0.01 | 0.03 | 1.00 | 39859 | 39731 |
| *sex: males* | *0.21* | *0.07* | *0.07* | *0.36* | *1.00* | *40228* | *39669* |
| sex: females and males | 0.33 | 0.22 | -0.08 | 0.78 | 1.00 | 40057 | 39923 |
| environment: wild | -0.15 | 0.24 | -0.64 | 0.30 | 1.00 | 40079 | 39805 |
| carotenoids: transformed | -0.12 | 0.18 | -0.49 | 0.23 | 1.00 | 39693 | 39431 |
| color: yellow | -0.10 | 0.18 | -0.45 | 0.27 | 1.00 | 38875 | 39581 |
| ornament: plumage | -0.06 | 0.28 | -0.62 | 0.50 | 1.00 | 40505 | 39417 |
| age: year1 | 0.03 | 0.11 | -0.19 | 0.24 | 1.00 | 38785 | 39591 |
| age: year1 and adults | -0.03 | 0.08 | -0.20 | 0.13 | 1.00 | 39064 | 38230 |
| **(B) Full model without phylogeny** |  |  |  |  |  |  |  |
| **Random terms** | **SD** | **Est.Error** | **l-95%CI** | **u-95%CI** | **Rhat** | **Bulk_ESS** | **Tail_ESS** |
| ~species | 0.15 | 0.09 | 0.01 | 0.35 | 1.00 | 39305 | 38622 |
| ~residual | 0.16 | 0.04 | 0.10 | 0.24 | 1.00 | 40475 | 39190 |
| ~phylogeny | NA | NA | NA | NA | NA | NA | NA |
| phylogenetic signal lambda | NA | NA | NA | NA |  |  |  |
| **Fixed effects** | **Estimate** | **Est.Error** | **l-95%CI** | **u-95%CI** | **Rhat** | **Bulk_ESS** | **Tail_ESS** |
| Intercept | 0.03 | 0.29 | -0.55 | 0.61 | 1.00 | 40022 | 39511 |
| maximum lifespan | 0.01 | 0.01 | -0.01 | 0.04 | 1.00 | 38932 | 39296 |
| monitoring years | 0.01 | 0.01 | -0.01 | 0.03 | 1.00 | 39420 | 39453 |
| *sex: males* | *0.21* | *0.07* | *0.07* | *0.35* | *1.00* | *39842* | *39981* |
| sex: females and males | 0.32 | 0.20 | -0.06 | 0.73 | 1.00 | 38984 | 36600 |
| environment: wild | -0.12 | 0.22 | -0.60 | 0.30 | 1.00 | 38102 | 38452 |
| carotenoids: transformed | -0.12 | 0.17 | -0.47 | 0.22 | 1.00 | 39702 | 39108 |
| color: yellow | -0.12 | 0.17 | -0.45 | 0.22 | 1.00 | 39723 | 39837 |
| ornament: plumage | -0.07 | 0.23 | -0.54 | 0.38 | 1.00 | 39528 | 39971 |
| age: year1 | 0.02 | 0.11 | -0.19 | 0.23 | 1.00 | 40131 | 37175 |
| age: year1 and adults | -0.04 | 0.08 | -0.20 | 0.12 | 1.00 | 39859 | 39471 |

**Table S3.** Final models (A) with and (B) without phylogeny. The levels in italics are statistically significant. Abbreviations are as in Table S2.

| **(A) Final model with phylogeny** |  |  |  |  |  |  |  |
| --- | --- | --- | --- | --- | --- | --- | --- |
| **Random terms** | **SD** | **Est.Error** | **l-95%CI** | **u-95%CI** | **Rhat** | **Bulk_ESS** | **Tail_ESS** |
| ~species | 0.06 | 0.05 | 0.00 | 0.19 | 1.00 | 40653 | 39846 |
| ~residual | 0.15 | 0.03 | 0.09 | 0.22 | 1.00 | 40435 | 39831 |
| ~phylogeny | 0.15 | 0.09 | 0.01 | 0.35 | 1.00 | 37592 | 37336 |
| phylogenetic signal lambda | 0.41 | 0.25 | 0.00 | 0.86 |  |  |  |
| **Fixed effects** | **Estimate** | **Est.Error** | **l-95%CI** | **u-95%CI** | **Rhat** | **Bulk_ESS** | **Tail_ESS** |
| Intercept females | -0.04 | 0.12 | -0.25 | 0.22 | 1.00 | 39450 | 39802 |
| *sex: males* | *0.20* | *0.06* | *0.07* | *0.32* | *1.00* | *40033* | *39845* |
| *sex: females and males* | *0.29* | *0.15* | *0.01* | *0.61* | *1.00* | *40136* | *40005* |
| **(B) Final model without phylogeny** |  |  |  |  |  |  |  |
| **Random terms** | **SD** | **Est.Error** | **l-95%CI** | **u-95%CI** | **Rhat** | **Bulk_ESS** | **Tail_ESS** |
| ~species | 0.07 | 0.06 | 0.00 | 0.21 | 1.00 | 40112 | 40045 |
| ~residual | 0.16 | 0.03 | 0.10 | 0.23 | 1.00 | 39626 | 39188 |
| ~phylogeny | NA | NA | NA | NA | NA | NA | NA |
| phylogenetic signal lambda | NA | NA | NA | NA |  |  |  |
| **Fixed effects** | **Estimate** | **Est.Error** | **l-95%CI** | **u-95%CI** | **Rhat** | **Bulk_ESS** | **Tail_ESS** |
| Intercept females | -0.10 | 0.05 | -0.21 | 0.01 | 1.00 | 39884 | 39549 |
| *sex: males* | *0.20* | *0.06* | *0.08* | *0.33* | *1.00* | *38415* | *39427* |
| *sex: females and males* | *0.28* | *0.13* | *0.04* | *0.56* | *1.00* | *39861* | *39930* |

**Table S4.**
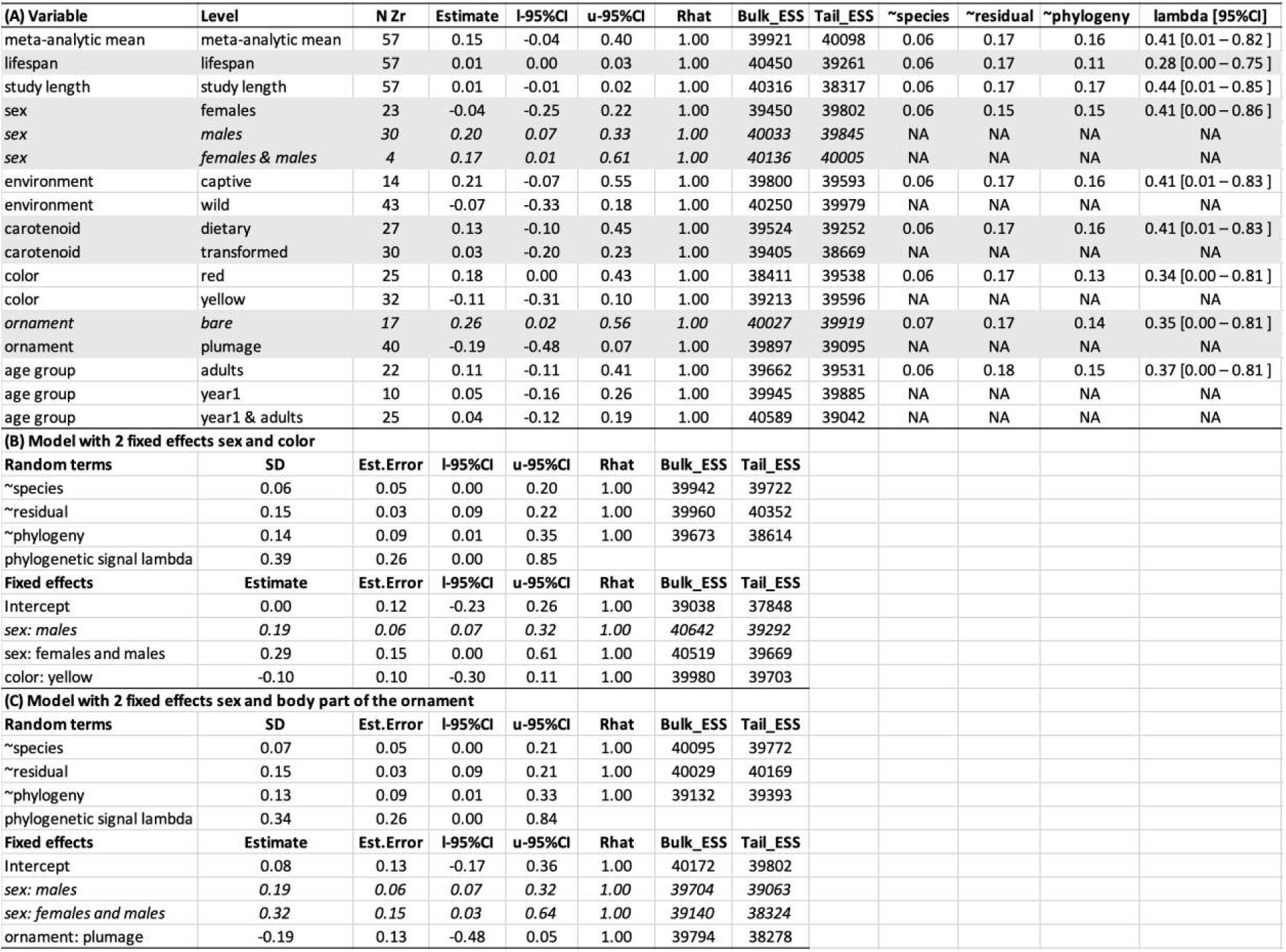
(A) Phylogenetic models per predictor variable used to identify the level-specific effect sizes for Figure 3. The shaded and white areas represent different variables. (B) Phylogenetic model with 2 variables sex and colour (red vs. yellow). (C) Phylogenetic model with 2 variables sex and ornament type (bare parts vs. plumage) The levels in italics are statistically significant. Abbreviations: N Zr = the number of effect sizes extracted from the literature per level; lambda = phylogenetic signal of the model. Other abbreviations are as in Table S2.

| (A) Variable | Level | N Zr | Estimate | I-95%CI | u-95%CI | Rhat | Bulk_ESS | Tail_ESS | ~species | ~residual | ~phylogeny | lambda [95%CI] |
| --- | --- | --- | --- | --- | --- | --- | --- | --- | --- | --- | --- | --- |
| meta-analytic mean | meta-analytic mean | 57 | 0.15 | -0.04 | 0.40 | 1.00 | 39921 | 40098 | 0.06 | 0.17 | 0.16 | 0.41 [0.01 – 0.82 ] |
| lifespan | lifespan | 57 | 0.01 | 0.00 | 0.03 | 1.00 | 40450 | 39261 | 0.06 | 0.17 | 0.11 | 0.28 [0.00 – 0.75 ] |
| study length | study length | 57 | 0.01 | -0.01 | 0.02 | 1.00 | 40316 | 38317 | 0.06 | 0.17 | 0.17 | 0.44 [0.01 – 0.85 ] |
| sex | females | 23 | -0.04 | -0.25 | 0.22 | 1.00 | 39450 | 39802 | 0.06 | 0.15 | 0.15 | 0.41 [0.00 – 0.86 ] |
| sex | *males* | 30 | 0.20 | 0.07 | 0.33 | 1.00 | 40033 | 39845 | NA | NA | NA | NA |
| sex | *females & males* | 4 | 0.17 | 0.01 | 0.61 | 1.00 | 40136 | 40005 | NA | NA | NA | NA |
| environment | captive | 14 | 0.21 | -0.07 | 0.55 | 1.00 | 39800 | 39593 | 0.06 | 0.17 | 0.16 | 0.41 [0.01 – 0.83 ] |
| environment | wild | 43 | -0.07 | -0.33 | 0.18 | 1.00 | 40250 | 39979 | NA | NA | NA | NA |
| carotenoid | dietary | 27 | 0.13 | -0.10 | 0.45 | 1.00 | 39524 | 39252 | 0.06 | 0.17 | 0.16 | 0.41 [0.01 – 0.83 ] |
| carotenoid | transformed | 30 | 0.03 | -0.20 | 0.23 | 1.00 | 39405 | 38669 | NA | NA | NA | NA |
| color | red | 25 | 0.18 | 0.00 | 0.43 | 1.00 | 38411 | 39538 | 0.06 | 0.17 | 0.13 | 0.34 [0.00 – 0.81 ] |
| color | yellow | 32 | -0.11 | -0.31 | 0.10 | 1.00 | 39213 | 39596 | NA | NA | NA | NA |
| ornament | *bare* | 17 | 0.26 | 0.02 | 0.56 | 1.00 | 40027 | 39919 | 0.07 | 0.17 | 0.14 | 0.35 [0.00 – 0.81 ] |
| ornament | plumage | 40 | -0.19 | -0.48 | 0.07 | 1.00 | 39897 | 39095 | NA | NA | NA | NA |
| age group | adults | 22 | 0.11 | -0.11 | 0.41 | 1.00 | 39662 | 39531 | 0.06 | 0.18 | 0.15 | 0.37 [0.00 – 0.81 ] |
| age group | year1 | 10 | 0.05 | -0.16 | 0.26 | 1.00 | 39945 | 39885 | NA | NA | NA | NA |
| age group | year1 & adults | 25 | 0.04 | -0.12 | 0.19 | 1.00 | 40589 | 39042 | NA | NA | NA | NA |
| **(B) Model with 2 fixed effects sex and color** |  |  |  |  |  |  |  |  |  |  |  |  |
| Random terms | SD | Est.Error | I-95%CI | u-95%CI | Rhat | Bulk_ESS | Tail_ESS |  |  |  |  |  |
| ~species | 0.06 | 0.05 | 0.00 | 0.20 | 1.00 | 39942 | 39722 |  |  |  |  |  |
| ~residual | 0.15 | 0.03 | 0.09 | 0.22 | 1.00 | 39960 | 40352 |  |  |  |  |  |
| ~phylogeny | 0.14 | 0.09 | 0.01 | 0.35 | 1.00 | 39673 | 38614 |  |  |  |  |  |
| phylogenetic signal lambda | 0.39 | 0.26 | 0.00 | 0.85 |  |  |  |  |  |  |  |  |
| Fixed effects | Estimate | Est.Error | I-95%CI | u-95%CI | Rhat | Bulk_ESS | Tail_ESS |  |  |  |  |  |
| Intercept | 0.00 | 0.12 | -0.23 | 0.26 | 1.00 | 39038 | 37848 |  |  |  |  |  |
| sex: *males* | 0.19 | 0.06 | 0.07 | 0.32 | 1.00 | 40642 | 39292 |  |  |  |  |  |
| sex: females and males | 0.29 | 0.15 | 0.00 | 0.61 | 1.00 | 40519 | 39669 |  |  |  |  |  |
| color: yellow | -0.10 | 0.10 | -0.30 | 0.11 | 1.00 | 39980 | 39703 |  |  |  |  |  |
| **(C) Model with 2 fixed effects sex and body part of the ornament** |  |  |  |  |  |  |  |  |  |  |  |  |
| Random terms | SD | Est.Error | I-95%CI | u-95%CI | Rhat | Bulk_ESS | Tail_ESS |  |  |  |  |  |
| ~species | 0.07 | 0.05 | 0.00 | 0.21 | 1.00 | 40095 | 39772 |  |  |  |  |  |
| ~residual | 0.15 | 0.03 | 0.09 | 0.21 | 1.00 | 40029 | 40169 |  |  |  |  |  |
| ~phylogeny | 0.13 | 0.09 | 0.01 | 0.33 | 1.00 | 39132 | 39393 |  |  |  |  |  |
| phylogenetic signal lambda | 0.34 | 0.26 | 0.00 | 0.84 |  |  |  |  |  |  |  |  |
| Fixed effects | Estimate | Est.Error | I-95%CI | u-95%CI | Rhat | Bulk_ESS | Tail_ESS |  |  |  |  |  |
| Intercept | 0.08 | 0.13 | -0.17 | 0.36 | 1.00 | 40172 | 39802 |  |  |  |  |  |
| sex: *males* | 0.19 | 0.06 | 0.07 | 0.32 | 1.00 | 39704 | 39063 |  |  |  |  |  |
| sex: females and males | 0.32 | 0.15 | 0.03 | 0.64 | 1.00 | 39140 | 38324 |  |  |  |  |  |
| ornament: plumage | -0.19 | 0.13 | -0.48 | 0.05 | 1.00 | 39794 | 38278 |  |  |  |  |  |

**Table S5.** Analyses of sex-specific effects for the variables carotenoids (dietary vs. transformed), colour (yellow vs. red) and ornament (bare parts vs. plumage). The mixed-sex group is excluded because it has too few data (4 effect sizes, see Table S1). Abbreviations are as in Table S2.

| **(A) Sex-specific interactions for a model with phylogeny** |  |  |  |  |  |  |  |
| --- | --- | --- | --- | --- | --- | --- | --- |
| **Random terms** | **SD** | **Est.Error** | **l-95%CI** | **u-95%CI** | **Rhat** | **Bulk_ESS** | **Tail_ESS** |
| ~species | 0.10 | 0.09 | 0.00 | 0.33 | 1.00 | 40288 | 39998 |
| ~residual | 0.17 | 0.04 | 0.10 | 0.25 | 1.00 | 40258 | 37616 |
| ~phylogeny | 0.16 | 0.13 | 0.01 | 0.48 | 1.00 | 40888 | 39938 |
| phylogenetic signal lambda | 0.34 | 0.27 | 0.00 | 0.87 |  |  |  |
| **Fixed effects** | **Estimate** | **Est.Error** | **l-95%CI** | **u-95%CI** | **Rhat** | **Bulk_ESS** | **Tail_ESS** |
| maximum lifespan | -0.01 | 0.02 | -0.05 | 0.02 | 1.00 | 40755 | 39756 |
| monitoring years | 0.01 | 0.01 | -0.01 | 0.03 | 1.00 | 40353 | 39101 |
| environment: wild | -0.02 | 0.25 | -0.52 | 0.45 | 1.00 | 40466 | 39373 |
| age: year1 | 0.05 | 0.12 | -0.17 | 0.28 | 1.00 | 39807 | 40009 |
| age: year1 and adults | 0.06 | 0.10 | -0.14 | 0.26 | 1.00 | 39934 | 39676 |
| females intercept | 0.03 | 0.41 | -0.78 | 0.82 | 1.00 | 38398 | 39336 |
| males intercept | 0.39 | 0.31 | -0.20 | 1.01 | 1.00 | 38254 | 39111 |
| females transformed | 0.00 | 0.28 | -0.54 | 0.56 | 1.00 | 37250 | 39017 |
| females yellow | -0.09 | 0.33 | -0.73 | 0.58 | 1.00 | 39133 | 39571 |
| females plumage | 0.05 | 0.34 | -0.64 | 0.71 | 1.00 | 39668 | 40009 |
| males transformed | -0.08 | 0.28 | -0.64 | 0.45 | 1.00 | 37911 | 39185 |
| males yellow | 0.08 | 0.32 | -0.55 | 0.71 | 1.00 | 39247 | 39930 |
| males plumage | -0.24 | 0.24 | -0.73 | 0.23 | 1.00 | 38484 | 39203 |
| **(B) Sex-specific interactions for a model without phylogeny** |  |  |  |  |  |  |  |
| **Random terms** | **SD** | **Est.Error** | **l-95%CI** | **u-95%CI** | **Rhat** | **Bulk_ESS** | **Tail_ESS** |
| ~species | 0.11 | 0.08 | 0.00 | 0.32 | 1.00 | 39973 | 39179 |
| ~residual | 0.18 | 0.04 | 0.11 | 0.26 | 1.00 | 39763 | 39656 |
| ~phylogeny | NA | NA | NA | NA | NA | NA | NA |
| phylogenetic signal lambda | NA | NA | NA | NA |  |  |  |
| **Fixed effects** | **Estimate** | **Est.Error** | **l-95%CI** | **u-95%CI** | **Rhat** | **Bulk_ESS** | **Tail_ESS** |
| maximum lifespan | -0.01 | 0.02 | -0.05 | 0.02 | 1.00 | 39821 | 39242 |
| monitoring years | 0.01 | 0.01 | -0.01 | 0.03 | 1.00 | 39227 | 38991 |
| environment: wild | -0.01 | 0.22 | -0.46 | 0.41 | 1.00 | 39751 | 39267 |
| age: year1 | 0.05 | 0.11 | -0.17 | 0.27 | 1.00 | 39693 | 39957 |
| age: year1 and adults | 0.06 | 0.10 | -0.14 | 0.25 | 1.00 | 40404 | 39395 |
| females intercept | 0.01 | 0.36 | -0.70 | 0.72 | 1.00 | 40505 | 39804 |
| males intercept | 0.40 | 0.31 | -0.20 | 1.01 | 1.00 | 39920 | 39619 |
| females transformed | 0.00 | 0.27 | -0.53 | 0.55 | 1.00 | 40361 | 39496 |
| females yellow | -0.12 | 0.32 | -0.74 | 0.51 | 1.00 | 39952 | 39763 |
| females plumage | 0.06 | 0.30 | -0.54 | 0.63 | 1.00 | 39249 | 39971 |
| males transformed | -0.07 | 0.28 | -0.62 | 0.48 | 1.00 | 39932 | 39477 |
| males yellow | 0.10 | 0.32 | -0.53 | 0.73 | 1.00 | 39862 | 39438 |
| males plumage | -0.29 | 0.25 | -0.78 | 0.19 | 1.00 | 39370 | 40228 |

**Table S6.**
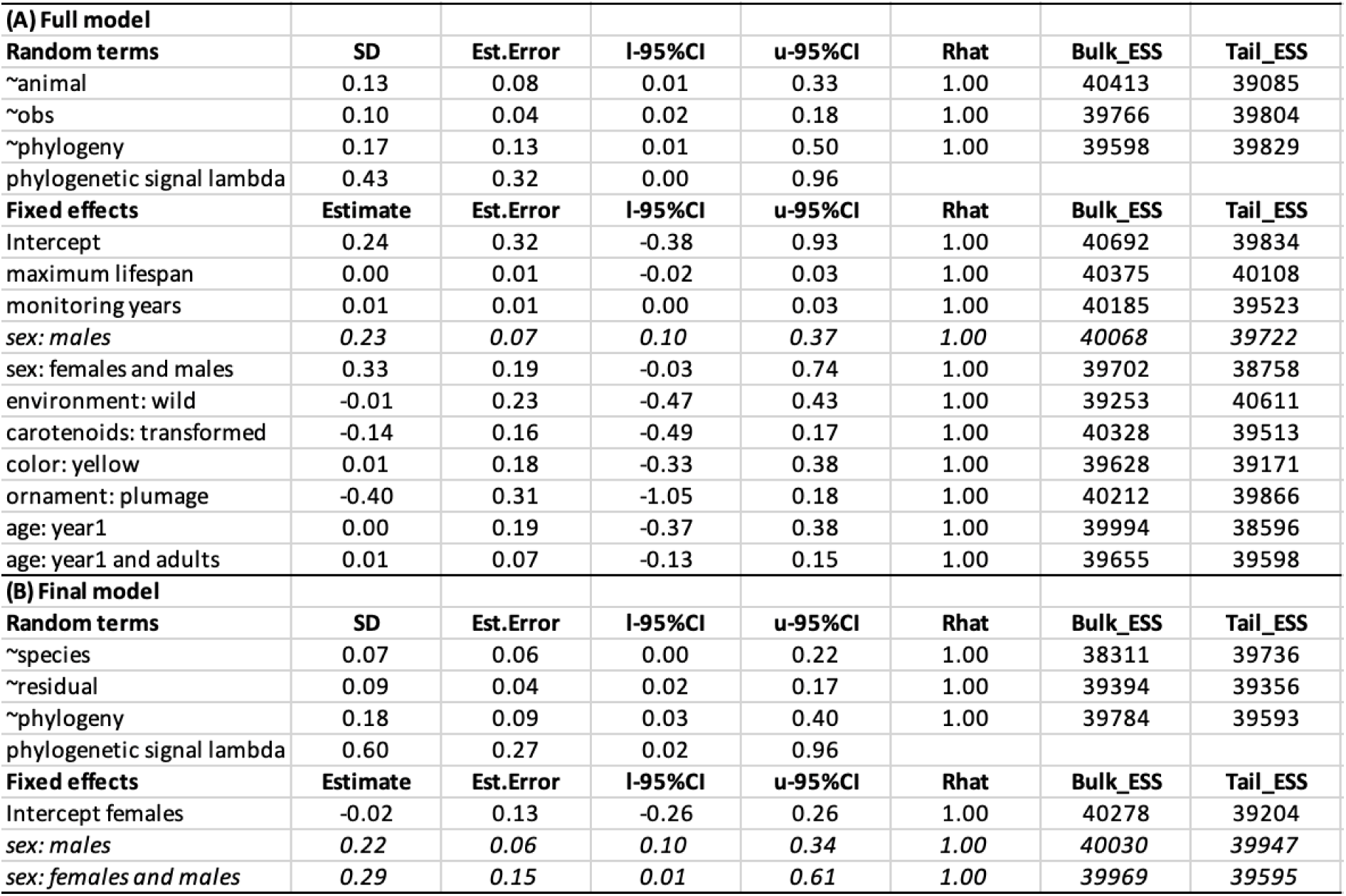
Full and final phylogenetic models for linear effect sizes using the data in Figure S3. The levels in italics are statistically significant. Abbreviations are as in Table S2.

**Table S7.** Final models (A) with and (B) without phylogeny for the 6 species with sex-specific data on both sexes separately. The 6 species are the Blue tit (*Cyanistes caeruleus*), Zebra finch (*Taeniopygia guttata*), House finch (*Haemorhous mexicanus*), Common yellowthroat (*Geothlypis trichas*), American redstart (*Setophaga ruticilla*), and White-throated sparrow (*Zonotrichia albicollis*). The levels in italics are statistically significant. Abbreviations are as in Table S2.

| **(A) Final model with phylogeny for the 6 species with data for both males and females separately** |  |  |  |  |  |  |  |
| --- | --- | --- | --- | --- | --- | --- | --- |
| **Random terms** | **SD** | **Est.Error** | **l-95%CI** | **u-95%CI** | **Rhat** | **Bulk_ESS** | **Tail_ESS** |
| ~species | 0.08 | 0.08 | 0.00 | 0.29 | 1.00 | 38773 | 36836 |
| ~residual | 0.17 | 0.04 | 0.09 | 0.25 | 1.00 | 39348 | 39710 |
| ~phylogeny | 0.12 | 0.12 | 0.00 | 0.44 | 1.00 | 38937 | 33360 |
| phylogenetic signal lambda | 0.25 | 0.25 | 0.00 | 0.84 |  |  |  |
| **Fixed effects** | **Estimate** | **Est.Error** | **l-95%CI** | **u-95%CI** | **Rhat** | **Bulk_ESS** | **Tail_ESS** |
| Intercept females | -0.11 | 0.13 | -0.38 | 0.16 | 1.00 | 37312 | 36070 |
| *sex: males* | *0.22* | *0.08* | *0.07* | *0.37* | *1.00* | *40162* | *39759* |
| **(B) Final model without phylogeny for the 6 species with data for both males and females separately** |  |  |  |  |  |  |  |
| **Random terms** | **SD** | **Est.Error** | **l-95%CI** | **u-95%CI** | **Rhat** | **Bulk_ESS** | **Tail_ESS** |
| ~species | 0.06 | 0.06 | 0.00 | 0.22 | 1.00 | 39412 | 39775 |
| ~residual | 0.16 | 0.04 | 0.09 | 0.25 | 1.00 | 40080 | 39717 |
| ~phylogeny | NA | NA | NA | NA | NA | NA | NA |
| phylogenetic signal lambda | NA | NA | NA | NA |  |  |  |
| **Fixed effects** | **Estimate** | **Est.Error** | **l-95%CI** | **u-95%CI** | **Rhat** | **Bulk_ESS** | **Tail_ESS** |
| Intercept females | -0.11 | 0.07 | -0.25 | 0.02 | 1.00 | 39993 | 39427 |
| *sex: males* | *0.22* | *0.08* | *0.08* | *0.37* | *1.00* | *39364* | *38534* |

**Table S8.**
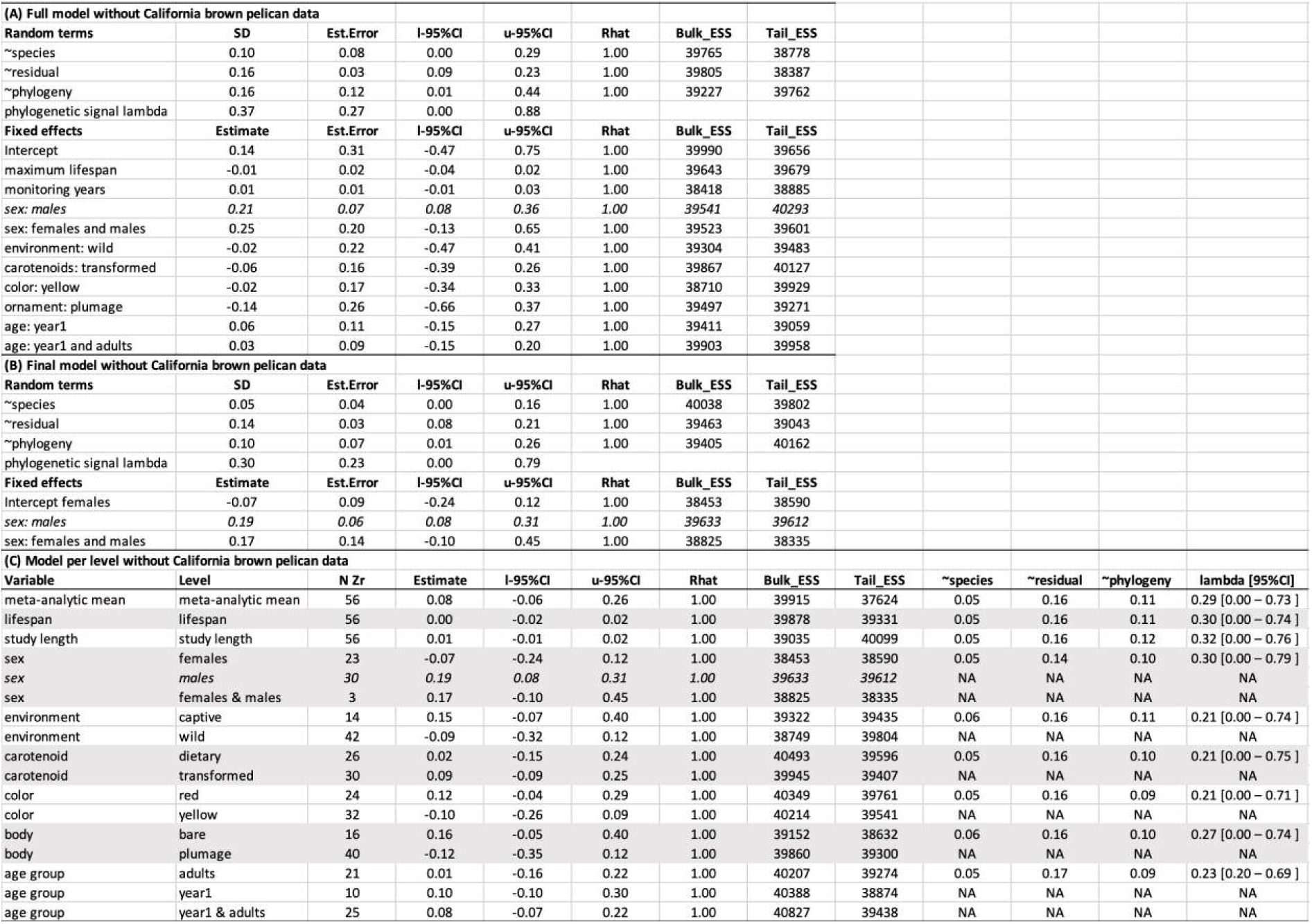
Full (A), final (B) and level-specific (C) models without California brown pelican (*Pelecanus occidentalis*) data. The levels in italics are statistically significant. The results in (C) are plotted in Figure S5. For (C), abbreviations are N Zr = the number of effect sizes extracted from the literature per level; lambda = phylogenetic signal of the model. Other abbreviations are as in Table S2.

| (A) Full model without California brown pelican data |  |  |  |  |  |  |  |  |  |  |  |  |
| --- | --- | --- | --- | --- | --- | --- | --- | --- | --- | --- | --- | --- |
| Random terms | SD | Est.Error | l-95%CI | u-95%CI | Rhat | Bulk_ESS | Tail_ESS |  |  |  |  |  |
| ~species | 0.10 | 0.08 | 0.00 | 0.29 | 1.00 | 39765 | 38778 |  |  |  |  |  |
| ~residual | 0.16 | 0.03 | 0.09 | 0.23 | 1.00 | 39805 | 38387 |  |  |  |  |  |
| ~phylogeny | 0.16 | 0.12 | 0.01 | 0.44 | 1.00 | 39227 | 39762 |  |  |  |  |  |
| phylogenetic signal lambda | 0.37 | 0.27 | 0.00 | 0.88 |  |  |  |  |  |  |  |  |
| Fixed effects | Estimate | Est.Error | l-95%CI | u-95%CI | Rhat | Bulk_ESS | Tail_ESS |  |  |  |  |  |
| Intercept | 0.14 | 0.31 | -0.47 | 0.75 | 1.00 | 39990 | 39656 |  |  |  |  |  |
| maximum lifespan | -0.01 | 0.02 | -0.04 | 0.02 | 1.00 | 39643 | 39679 |  |  |  |  |  |
| monitoring years | 0.01 | 0.01 | -0.01 | 0.03 | 1.00 | 38418 | 38885 |  |  |  |  |  |
| sex: males | 0.21 | 0.07 | 0.08 | 0.36 | 1.00 | 39541 | 40293 |  |  |  |  |  |
| sex: females and males | 0.25 | 0.20 | -0.13 | 0.65 | 1.00 | 39523 | 39601 |  |  |  |  |  |
| environment: wild | -0.02 | 0.22 | -0.47 | 0.41 | 1.00 | 39304 | 39483 |  |  |  |  |  |
| carotenoids: transformed | -0.06 | 0.16 | -0.39 | 0.26 | 1.00 | 39867 | 40127 |  |  |  |  |  |
| color: yellow | -0.02 | 0.17 | -0.34 | 0.33 | 1.00 | 38710 | 39929 |  |  |  |  |  |
| ornament: plumage | -0.14 | 0.26 | -0.66 | 0.37 | 1.00 | 39497 | 39271 |  |  |  |  |  |
| age: year1 | 0.06 | 0.11 | -0.15 | 0.27 | 1.00 | 39411 | 39059 |  |  |  |  |  |
| age: year1 and adults | 0.03 | 0.09 | -0.15 | 0.20 | 1.00 | 39903 | 39958 |  |  |  |  |  |
| (B) Final model without California brown pelican data |  |  |  |  |  |  |  |  |  |  |  |  |
| Random terms | SD | Est.Error | l-95%CI | u-95%CI | Rhat | Bulk_ESS | Tail_ESS |  |  |  |  |  |
| ~species | 0.05 | 0.04 | 0.00 | 0.16 | 1.00 | 40038 | 39802 |  |  |  |  |  |
| ~residual | 0.14 | 0.03 | 0.08 | 0.21 | 1.00 | 39463 | 39043 |  |  |  |  |  |
| ~phylogeny | 0.10 | 0.07 | 0.01 | 0.26 | 1.00 | 39405 | 40162 |  |  |  |  |  |
| phylogenetic signal lambda | 0.30 | 0.23 | 0.00 | 0.79 |  |  |  |  |  |  |  |  |
| Fixed effects | Estimate | Est.Error | l-95%CI | u-95%CI | Rhat | Bulk_ESS | Tail_ESS |  |  |  |  |  |
| Intercept females | -0.07 | 0.09 | -0.24 | 0.12 | 1.00 | 38453 | 38590 |  |  |  |  |  |
| sex: males | 0.19 | 0.06 | 0.08 | 0.31 | 1.00 | 39633 | 39612 |  |  |  |  |  |
| sex: females and males | 0.17 | 0.14 | -0.10 | 0.45 | 1.00 | 38825 | 38335 |  |  |  |  |  |
| (C) Model per level without California brown pelican data |  |  |  |  |  |  |  |  |  |  |  |  |
| Variable | Level | N Zr | Estimate | l-95%CI | u-95%CI | Rhat | Bulk_ESS | Tail_ESS | ~species | ~residual | ~phylogeny | lambda [95%CI] |
| meta-analytic mean | meta-analytic mean | 56 | 0.08 | -0.06 | 0.26 | 1.00 | 39915 | 37624 | 0.05 | 0.16 | 0.11 | 0.29 [0.00 – 0.73] |
| lifespan | lifespan | 56 | 0.00 | -0.02 | 0.02 | 1.00 | 39878 | 39331 | 0.05 | 0.16 | 0.11 | 0.30 [0.00 – 0.74] |
| study length | study length | 56 | 0.01 | -0.01 | 0.02 | 1.00 | 39035 | 40099 | 0.05 | 0.16 | 0.12 | 0.32 [0.00 – 0.76] |
| sex | females | 23 | -0.07 | -0.24 | 0.12 | 1.00 | 38453 | 38590 | 0.05 | 0.14 | 0.10 | 0.30 [0.00 – 0.79] |
| sex | males | 30 | 0.19 | 0.08 | 0.31 | 1.00 | 39633 | 39612 | NA | NA | NA | NA |
| sex | females & males | 3 | 0.17 | -0.10 | 0.45 | 1.00 | 38825 | 38335 | NA | NA | NA | NA |
| environment | captive | 14 | 0.15 | -0.07 | 0.40 | 1.00 | 39322 | 39435 | 0.06 | 0.16 | 0.11 | 0.21 [0.00 – 0.74] |
| environment | wild | 42 | -0.09 | -0.32 | 0.12 | 1.00 | 38749 | 39804 | NA | NA | NA | NA |
| carotenoid | dietary | 26 | 0.02 | -0.15 | 0.24 | 1.00 | 40493 | 39596 | 0.05 | 0.16 | 0.10 | 0.21 [0.00 – 0.75] |
| carotenoid | transformed | 30 | 0.09 | -0.09 | 0.25 | 1.00 | 39945 | 39407 | NA | NA | NA | NA |
| color | red | 24 | 0.12 | -0.04 | 0.29 | 1.00 | 40349 | 39761 | 0.05 | 0.16 | 0.09 | 0.21 [0.00 – 0.71] |
| color | yellow | 32 | -0.10 | -0.26 | 0.09 | 1.00 | 40214 | 39541 | NA | NA | NA | NA |
| body | bare | 16 | 0.16 | -0.05 | 0.40 | 1.00 | 39152 | 38632 | 0.06 | 0.16 | 0.10 | 0.27 [0.00 – 0.74] |
| body | plumage | 40 | -0.12 | -0.35 | 0.12 | 1.00 | 39860 | 39300 | NA | NA | NA | NA |
| age group | adults | 21 | 0.01 | -0.16 | 0.22 | 1.00 | 40207 | 39274 | 0.05 | 0.17 | 0.09 | 0.23 [0.20 – 0.69] |
| age group | year1 | 10 | 0.10 | -0.10 | 0.30 | 1.00 | 40388 | 38874 | NA | NA | NA | NA |
| age group | year1 & adults | 25 | 0.08 | -0.07 | 0.22 | 1.00 | 40827 | 39438 | NA | NA | NA | NA |

**Table S9.** Final (A) and full (B) models with the phylogeny by McTavish et al. 2025.

| **(A) Full model with phylogeny by McTavish et al. 2025** |  |  |  |  |  |  |  |
| --- | --- | --- | --- | --- | --- | --- | --- |
| **Random terms** | **SD** | **Est.Error** | **l-95%CI** | **u-95%CI** | **Rhat** | **Bulk_ESS** | **Tail_ESS** |
| ~species | 0.12 | 0.09 | 0.01 | 0.33 | 1.00 | 40259 | 38614 |
| ~residual | 0.16 | 0.04 | 0.09 | 0.23 | 1.00 | 39648 | 38815 |
| ~phylogeny | 0.03 | 0.02 | 0.00 | 0.07 | 1.00 | 39197 | 38258 |
| phylogenetic signal lambda | 0.03 | 0.04 | 0.00 | 0.14 |  |  |  |
| **Fixed effects** | **Estimate** | **Est.Error** | **l-95%CI** | **u-95%CI** | **Rhat** | **Bulk_ESS** | **Tail_ESS** |
| Intercept | 0.09 | 0.36 | -0.64 | 0.84 | 1.00 | 39519 | 39290 |
| maximum lifespan | 0.01 | 0.01 | -0.02 | 0.04 | 1.00 | 40488 | 39533 |
| monitoring years | 0.01 | 0.01 | -0.01 | 0.03 | 1.00 | 39045 | 39926 |
| sex: *males* | 0.21 | 0.07 | 0.07 | 0.35 | 1.00 | 40102 | 38534 |
| sex: females and males | 0.31 | 0.23 | -0.13 | 0.80 | 1.00 | 39177 | 39972 |
| environment: wild | -0.14 | 0.23 | -0.62 | 0.30 | 1.00 | 40069 | 38842 |
| carotenoids: transformed | -0.12 | 0.18 | -0.48 | 0.22 | 1.00 | 39852 | 40243 |
| color: yellow | -0.09 | 0.17 | -0.43 | 0.26 | 1.00 | 39415 | 39483 |
| ornament: plumage | -0.02 | 0.26 | -0.55 | 0.51 | 1.00 | 40960 | 40352 |
| age: year1 | 0.02 | 0.11 | -0.19 | 0.24 | 1.00 | 38510 | 39392 |
| age: year1 and adults | -0.03 | 0.08 | -0.20 | 0.14 | 1.00 | 39692 | 39110 |
| **(B) Final model with phylogeny by McTavish et al. 2025** |  |  |  |  |  |  |  |
| **Random terms** | **SD** | **Est.Error** | **l-95%CI** | **u-95%CI** | **Rhat** | **Bulk_ESS** | **Tail_ESS** |
| ~species | 0.05 | 0.05 | 0.00 | 0.17 | 1.00 | 40040 | 39720 |
| ~residual | 0.14 | 0.03 | 0.08 | 0.21 | 1.00 | 39366 | 39490 |
| ~phylogeny | 0.02 | 0.01 | 0.00 | 0.05 | 1.00 | 40342 | 39680 |
| phylogenetic signal lambda | 0.02 | 0.03 | 0.10 |  |  |  |  |
| **Fixed effects** | **Estimate** | **Est.Error** | **l-95%CI** | **u-95%CI** | **Rhat** | **Bulk_ESS** | **Tail_ESS** |
| Intercept females | 0.04 | 0.16 | -0.25 | 0.38 | 1.00 | 39152 | 39040 |
| sex: *males* | 0.19 | 0.06 | 0.07 | 0.31 | 1.00 | 40214 | 38728 |
| sex: females and males | 0.27 | 0.16 | -0.03 | 0.60 | 1.00 | 39472 | 39719 |

## Notes

### Competing Interest Statement

The authors have declared no competing interest.

